# The *mex*-3 3’ untranslated region is essential for reproduction during temperature stress

**DOI:** 10.1101/2024.04.01.587367

**Authors:** Hannah E. Brown, Haik V. Varderesian, Sara A. Keane, Sean P. Ryder

## Abstract

Organisms must sense temperature and modify their physiology to survive environmental stress. Elevated temperature reduces fertility in most sexually reproducing organisms. Maternally supplied mRNAs are required for embryogenesis. They encode proteins that govern early embryonic patterning. RNA-binding proteins (RBPs) are major effectors of maternal mRNA regulation. MEX-3 is a conserved RBP essential for anterior patterning of *Caenorhabditis elegans* embryos. We previously demonstrated that the *mex-3* 3’ untranslated region (3’UTR) represses MEX-3 abundance in the germline yet is mostly dispensable for fertility. Here, we show that the 3’UTR is essential during thermal stress. Deletion of the 3’UTR causes a highly penetrant temperature sensitive embryonic lethality phenotype distinct from a *mex-3* null. Loss of the 3’UTR decreases MEX-3 abundance specifically in maturing oocytes and early embryos during temperature stress. Dysregulation of *mex-3* reprograms the thermal stress response by reducing the expression of hundreds of heat shock genes. We propose that the primary role of the *mex-3* 3’UTR is to buffer MEX-3 expression during fluctuating temperature, ensuring the robustness of oocyte maturation and embryogenesis.

## INTRODUCTION

Thermoregulation is critical for sexual reproduction throughout the plant and animal kingdoms (Abram et al., 2017; Hansen, 2009; Walsh et al., 2019). Mammals evolved external testicles to ensure that spermatogenesis occurs below body temperature (Lovegrove, 2014). In chickens and mice, short exposure to thermal stress results in prolonged reduction of sperm concentration (Walsh et al., 2019). In cows, ovulation and implantation are compromised in warmer seasons (Hansen, 2009). In humans, temperature stress can impact gestational health and developmental consequences in the children of affected mothers (Dreier et al., 2014).

Ectotherms, which do not thermoregulate, employ a variety of adaptations to ensure robust embryogenesis in thermal stress (Abram et al., 2017). Hornets regulate hive temperature by beating their wings at the nest entrance to ensure adequate ventilation (Riabinin et al., 2004). Turtle embryos reorient inside of the shell to ensure optimal temperature (Du et al., 2011). In certain reptiles and fish, sex determination is controlled by external temperature cues (Bull and Vogt, 1979; Conover and Heins, 1987). Although it is generally accepted that thermoregulation is fundamental to reproductive success, the molecular mechanisms that protect gametes and embryos from thermal stress are not well understood.

Several aspects of *Caenorhabditis elegans* reproduction are sensitive to temperature. The optimal reproductive temperature of this species is 20°C (Stiernagle, 2006). Prolonged thermal stress above 28°C prevents embryonic development and hatching (Aprison and Ruvinsky, 2014). Thermal stress above 31°C prevents egg production (Aprison and Ruvinsky, 2014). Prolonged exposure to 27°C causes transgenerational sterilization—the progeny of stressed adults are sterile (Begasse et al., 2015). L1 larva that experience temperature stress enter an alternative life cycle, producing temperature resistant dauer larva that can persist for long periods of time before re-entering the path to adulthood when conditions are more favorable (Byerly et al., 1976).

In general, the higher the temperature and the longer the exposure, the more significant the impact to reproductive fecundity. Yet even limited duration exposure to elevated temperature can affect gametogenesis, ovulation rate, and hatch rate, depending on time and temperature, with variable recovery rates upon shifting to the optimal environment (Anderson et al., 2011; Aprison and Ruvinsky, 2014; Begasse et al., 2015; Kurhanewicz et al., 2020; Rogers and Phillips, 2020). Short duration exposure to heat alters the sex ratio in *C. elegans*, increasing the number of chromosomal nondisjunction events that occur during spermatogenesis resulting in more XO males (as opposed to XX hermaphrodites) (Byerly et al., 1976). One of the earliest embryonic patterning events, the timing of the first cell division cycle, is sensitive to temperature (Begasse et al., 2015). The rate of the first cellular division scales exponentially with increasing temperature. The underlying mechanisms behind these phenomena are not clear, but the types of defects and their timing during the reproductive cycle suggest an important role for maternal gene products.

In most animals, maternally deposited proteins and mRNAs are critical to fertility and embryo development after fertilization (Farley and Ryder, 2008). Stored maternal mRNAs are regulated in *cis* by elements found within their untranslated regions and in *trans* by RNA-binding proteins (RBPs) and microRNAs (miRNAs). In *C. elegans*, the germline undergoes several important transitions, including a switch from mitosis to meiosis, a transition from individual germ cells to a syncytial state, differentiation into either sperm or eggs, and the maturation of oocytes (Albarqi and Ryder, 2022). Following fertilization, embryos undergo transitions that govern body axis formation, segregation of the germline from somatic cells, and cell fate specification prior to zygotic gene activation. Maternal mRNAs and RBPs contribute to each of these transitions in the germline and embryo.

MEX-3 is a conserved KH-domain RNA-binding protein required for anterior cell fate specification in early embryogenesis (**Fig. 1**) (Draper et al., 1996; Huang et al., 2002). Loss of function *mex-3* alleles cause anterior cells to adopt posterior like fates, leading to excess muscle and germline lineage cells in terminal embryos. In the adult germline, MEX-3 expression is restricted to the distal mitotic region and the proximal oocytes (Draper et al., 1996). In the embryo, expression is enriched in the anterior until the 8-16 cells stage, when expression is lost. In the posterior, the remaining MEX-3 protein is sequestered in P granules in germline progenitor cells (Draper et al., 1996). In animals depleted of sperm, MEX-3 aggregates into large RNP granules in oocytes (Elaswad et al., 2022; Jud et al., 2008).

**Figure 1.**
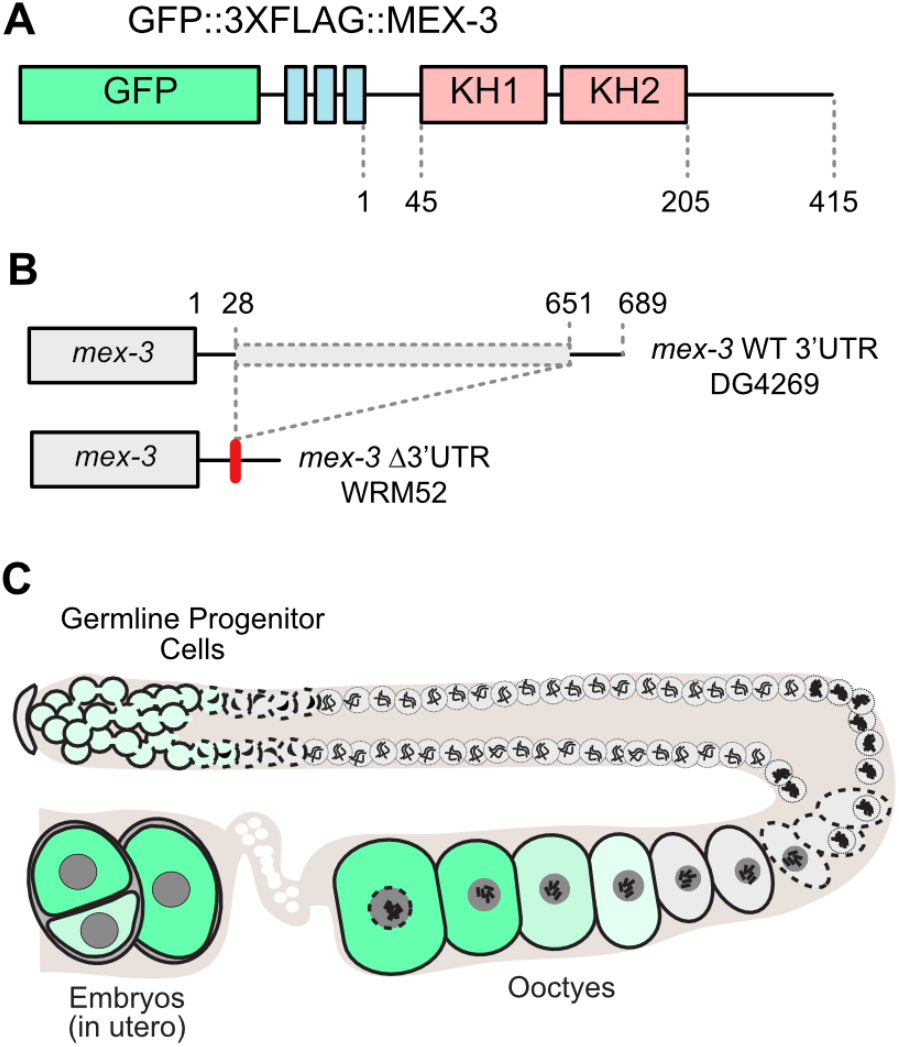
MEX-3 is a germline RBP required for anterior cell fate specification in the embryo. **A**. The endogenous MEX-3 gene contains two KH domains and was tagged with GFP and 3X FLAG at the N-terminus (DG4269) (Tsukamoto et al., 2017). **B**. The deletion allele that removes 624 of the 689 nucleotides of the 3’UTR was made in this strain background (WRM52) (Albarqi and Ryder, 2021). **C**. The germline MEX-3 expression is diagrammed in green, with darker colors indicating stronger expression.

The *mex-3* 3’UTR is sufficient to confer the MEX-3 expression pattern to a reporter transgene in the germline (Kaymak et al., 2016; Merritt et al., 2008). The 3’UTR is also necessary for patterned expression from the endogenous locus (Albarqi and Ryder, 2021). We previously showed that a 624 base pair deletion mutant in the endogenous 3’UTR (*mex-3 (spr9[*tn1753])*, hereafter referred to as *gfp::3xflag*::*mex-3*::Δ3’UTR) increases MEX-3 abundance throughout the germline. Despite extensive dysregulation, this mutant displays a modest reduction in total and viable brood (Albarqi and Ryder, 2021). Given the importance of MEX-3 to embryonic cell fate and the clear role of its 3’UTR to patterning its expression, we were surprised that loss of the 3’UTR did not yield a more striking reproductive phenotype. In this study, we tested the effect of stress on animals that lack the *mex-3* 3’UTR. Our results reveal that the 3’UTR is essential during thermal stress, and that animals harboring this mutation display a highly penetrant embryonic lethality phenotype that is distinct from the null allele. Our results suggest a primary role of the *mex-3* 3’UTR is to buffer MEX-3 protein expression during periods of thermal stress.

## RESULTS

### The mex-3 3’UTR is essential at elevated temperature

To determine if the *mex-3* 3’UTR is important to reproductive fecundity during stress, we recorded the brood size (total number of embryos produced) and hatch rate (viable progeny / total progeny) of 3’UTR deletion mutants as compared to a background matched control under a variety of environmental stress conditions. Animals were cultured at reduced and elevated temperature (15°C and 25°C), increased osmotic stress (350 mM NaCl), and pathogenic stress induced by feeding on *Pseudomonas aeruginosa* (PA14). We compared the results to animals grown under standard laboratory conditions (20°C, 50 mM NaCl, *Escherichia coli* (OP50); **Fig. 2**).

**Figure 2.**
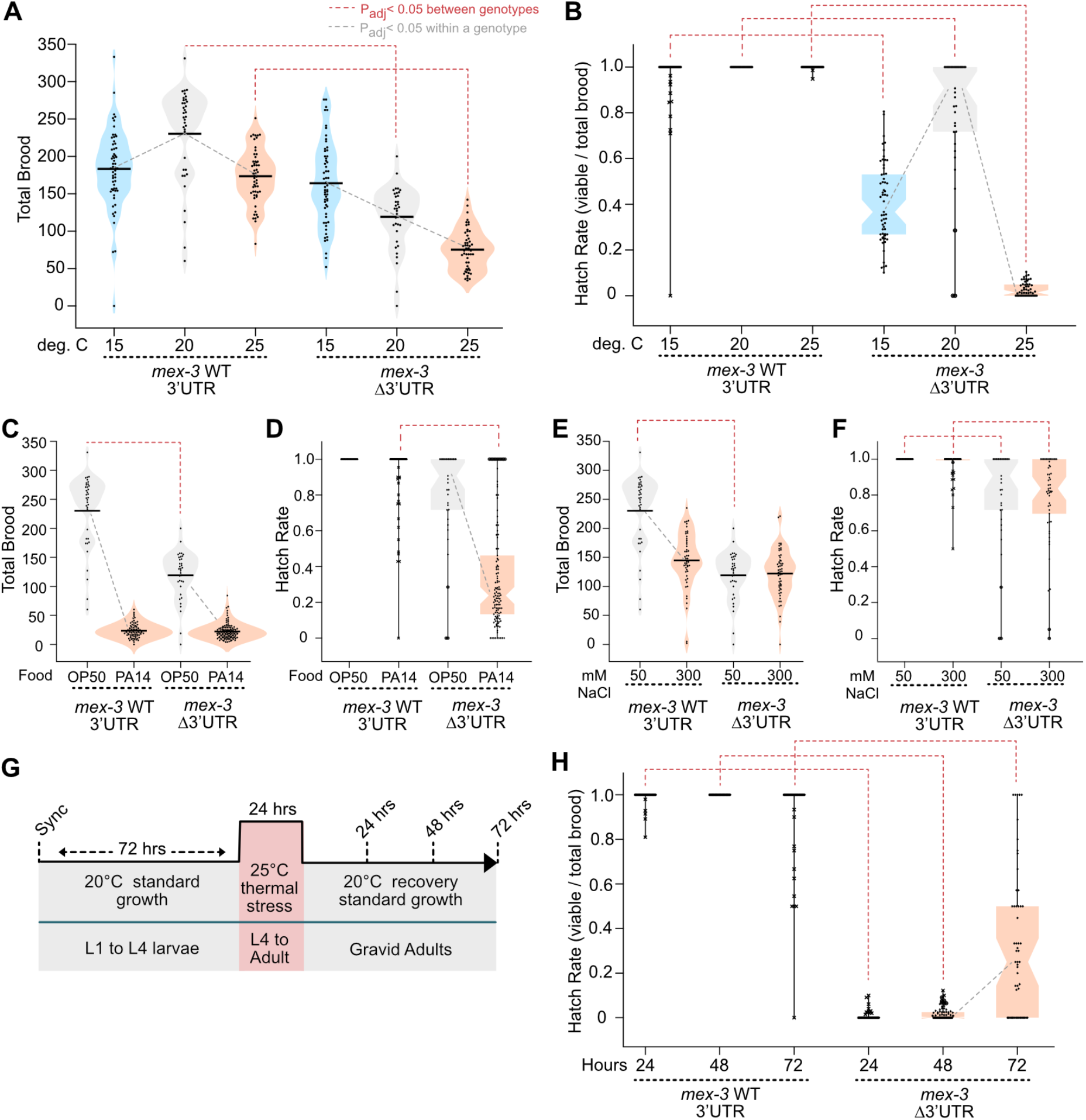
The *mex-3* 3’UTR is essential during temperature stress. **A**. Violin plots of the total brood for control and *mex-3* 3’UTR deletion strains. Blue, gray, and orange violins correspond to 15, 20, and 25 ºC growth. In all panels, gray lines indicate statistical significance within a genotype and red brackets indicates the same between genotypes in a one-way ANOVA with Bonferroni correction (P_adj_< 0.05). **B**. Box and whisker plots depicting the hatch rate of embryos from panel A. **C-F**. Brood size and hatch rate data for infectious stress (C-D) and osmotic stress (E-F). In C-D, the orange color indicates *Pseudomonas aeruginosa (PA14)* while gray indicates *E. coli* food. In E-F, the orange color indicates high salt (300 mM NaCl) while gray indicates standard salt (50 mM NaCl). **G**. Time course to assess reversibility of the temperature sensitive embryonic lethality phenotype. **H**. Box and whiskers plot of the hatch rate post recovery in each genotype.

The temperature dependence of the brood size produced by *C. elegans* Bristol N2 worms is bell shaped with a maximum near 18.2°C (Begasse et al., 2015). We observe a similar pattern with the wild-type *gfp::3xflag*::*mex-3*::WT 3’UTR control strain (DG4269; Mean Brood_WT_ = 183 ± 53 at 15°C, 230 ± 63 at 20°C, and 173 ± 36 at 25°C, P_adj_ <0.001, **Fig. 2A**). By contrast, the 3’UTR deletion strain shows a linear decrease in the number of embryos produced as temperature increases, with the maximum produced at 15°C (WRM52; Mean Brood_Δ3’UTR_ = 164 ± 55 at 15°C, 119 ± 42 at 20°C, and 75 ± 27 at 25°C, P_adj_ <0.001, **Fig. 2A**). At 15°C, both strains produce similar size broods (P_adj_ = 0.36), while at 25°C, the 3’UTR deletion mutant produces less than half of the control (Brood _Δ3’UTR-25_ / Brood _WT-25_ = 0.43, P_adj_ < 0.001). As such, increasing the temperature enhances the reduced fecundity phenotype nearly 2-fold, while lowering the temperature rescues the phenotype.

Changing the temperature also has a strong impact on the number of embryos that hatch in the *mex-3* 3’UTR deletion mutant (**Fig. 2B**). Consistent with our previous report, the 3’UTR deletion induces a modest reduction in the number of embryos that hatch compared to a background matched control under standard lab conditions (Hatch Rate_Δ3’UTR_ = 81%, P_adj_ = 0.00158) (Albarqi and Ryder, 2021). At 15°C, only 40% of the embryos hatch (P_adj_ < 0.001), and at 25°C, just 3% of the embryos hatch (P_adj_ < 0.001). By contrast, nearly all embryos produced by the control strain hatched at all three temperatures (Hatch Rate_WT_ = 96% at 15°C and 100% at 20°C and 25°C). The data show that the 3’UTR mutation induces a strong and highly penetrant embryonic lethality phenotype at 25°C, and a less penetrant embryonic lethality phenotype at 15°C. These phenotypes are distinct from the effect on total brood size described above.

Culturing the worms on the bacterial pathogen *Pseudomonas aeruginosa* (strain PA14) decreased the brood size of both strains by 90% (Mean Brood_WT_ = 26 ± 12 on PA14, Mean Brood_Δ3’UTR_ = 22 ± 13, P_adj_ = 1.0, **Fig. 2C**). However, 93% of the WT UTR embryos hatched, while only 35% of the Δ3’UTR mutant embryos hatched (P_adj_ < 0.001, **Fig. 2D**). As such, the 3’UTR mutation also causes a partially penetrant embryonic lethality phenotype under pathogenic stress.

Exposure to elevated sodium chloride reduced the total brood produced by both strains to a lesser extent than pathogenic stress, but there was no significant difference between genotypes (Fold Effect WT_300mM/50mM_ = 0.63, P_adj_ = <0.001; Fold Effect Δ3’UTR_300mM/50mM_ = 0.75, P_adj_ = <0.001, Fold Effect 300 mM NaCl_Δ3’UTR/WT_ = 0.85, P_adj_ = 0.47, **Fig. 2E**). Unlike thermal and pathogenic stress, exposure to high salt did not enhance the moderate embryonic lethality phenotype induced by the 3’UTR deletion mutant. (Fold Effect Δ3’UTR_300mM/50mM_ = 0.98, P_adj_ = <0.001, **Fig. 2F**). Together, the data show that some, but not all, stress conditions enhance the phenotype of the *mex-3* Δ3’UTR mutant.

The temperature sensitive embryonic lethality phenotype is only observed when the majority of the 3’UTR is deleted. Three shorter 3’UTR deletion alleles—Δ26–167, Δ328–517, and Δ515–648—had similar brood sizes and hatch rates to the wild type 3’UTR control (**Supplemental Fig. 1**).

### The temperature-induced phenotype is reversible

MEX-3 is expressed in germline progenitor cells, in oocytes, and early-stage embryos. It plays a role in maintaining progenitor cell totipotency in the germline and it is required for anterior cell fate specification in early embryos (Ariz et al., 2009; Ciosk et al., 2004; Draper et al., 1996). Heat stress could affect one or both processes. We reasoned that if heat stress modifies MEX-3 function in embryos, it might be rapidly reversible upon removal from stress because new embryos are produced approximately every 20 minutes. By contrast, if heat stress impacts MEX-3’s function in the germline, recovery might take more time, or affected animals may never recover.

To test these hypotheses, we isolated individual synchronized L1 larval stage worms and allowed them to mature for 72 hours under standard conditions (20°C, **Fig. 2G**). Then, we stressed the worms at 25°C for 24 hours during the late L4 to young adult stage when oogenesis begins. Worms were allowed to recover at 20°C for 24, 48, and 72 hours before measuring the hatch rate of the brood deposited on each plate. Treated animals were transferred to a new plate each day so we could monitor recovery as a function of time post stress. Our data show that the *mex-3* 3’UTR deletion mutant produced embryos with a hatch rate of 0.6% at 24 hours, 1.7% at 48 hours, and 33% at 72 hours post thermal stress (**Fig 2H**), showing partial recovery of fecundity in the window between 48 and 72 hours. The WT 3’UTR control strain had a hatch rate of 99%, 100%, and 94% across the same treatment regime and time span (P_adj_< 0.001). Our data show that while recovery of embryonic viability is possible, it takes between two and three days post thermal stress to restore viability, and recovery is not complete. Therefore, thermal stress likely affects some developmental process that requires days to recover.

### MEX-3 abundance is reduced in late-stage oocytes and early embryos during thermal stress

As shown in **Figure 1**, GFP::MEX-3 is expressed at low levels in the distal end of the germline and then disappears upon entry into meiosis. Expression resumes during oogenesis peaking in the most proximal oocyte undergoing maturation. Expression increases in early embryos before the protein disappears by the 16-32 cell stage. We previously showed that deletion of the *mex-3* 3’UTR causes a global increase in GFP::MEX-3 protein throughout the germline, revealing that the 3’UTR acts to repress MEX-3 expression (Albarqi and Ryder, 2021). To determine if the expression pattern is affected by temperature, we cultured control and 3’UTR deletion mutant animals at 25°C and collected images of young adult worms. The germline expression pattern of control animals does not appear to be affected by elevated temperature. Expression remains elevated throughout the germline at both temperatures in the 3’UTR deletion mutant (**Fig. 3A**).

**Figure 3.**
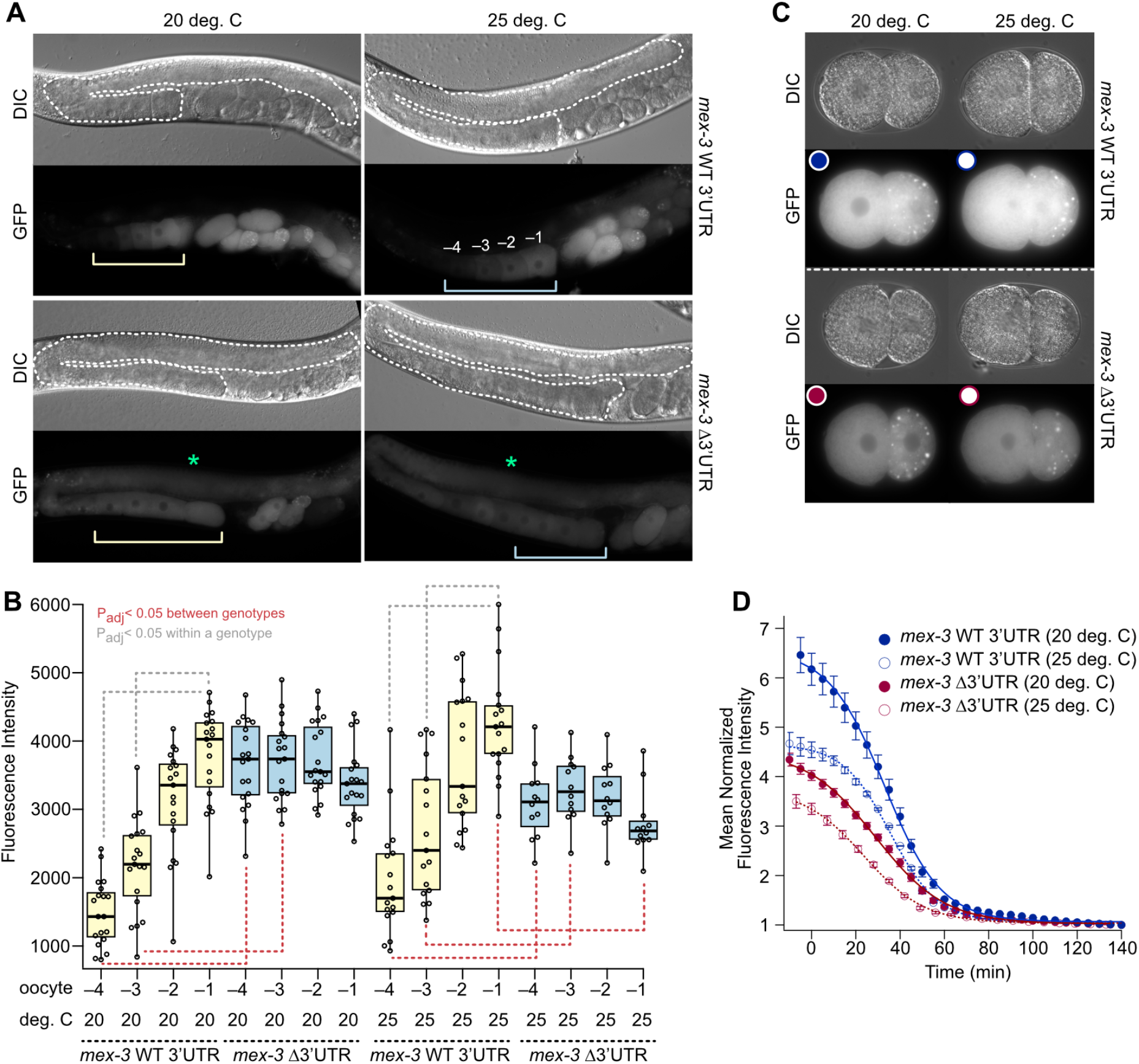
Elevated temperature decreases GFP::MEX-3 in maturing oocytes and embryos. **A**. DIC and GFP images of the control and *mex-3* 3’UTR deletion mutant strains. The dashed white line outlines a gonad, the green asterisk marks the syncitial region. The yellow and blue brackets mark developing oocytes. The -4 to -1 oocytes are labeled. **B**. Box and whisker plots of oocyte fluorescence at standard and elevated temperature. Statistical significance is indicated as in Figure 2. **C**. Still frames from embrogenesis movies at t=0. The colored circles correspond to the markers in panel D. **D**. Quantitation of GFP::MEX-3 in embryogenesis movies. The markers indicate the mean per time point of multiple movies per sample (n=2-5). The error bars are the standard error, and the curves are fits to a sigmoidal decay equation to determine the maximal fluorescence, the half-life, and the rate of decay.

Next, we quantitated GFP::MEX-3 abundance in oocytes by measuring mean pixel intensity in the four most proximal oocytes (−4 to -1, **Fig. 3A**). Control animals display the expected increase in GFP::MEX-3 abundance in older oocytes at both temperatures (**Fig. 3A-B**). In contrast, the expression of GFP::MEX-3 remains constant in the 3’UTR deletion mutant during oocyte development. At 25 ºC, the abundance of GFP::MEX-3 in the most proximal oocyte of the mutant is 1.5-fold less than controls. These results suggest that the 3’UTR contributes to increasing MEX-3 expression observed during oocyte maturation. They also suggest that MEX-3 insufficiency in oocytes contributes to the temperature sensitive phenotype.

To measure GFP::MEX-3 expression in embryos, we filmed embryogenesis and monitored the total fluorescence and the rate of decay as a function of time (**Supplemental Movies 1-2**). To facilitate comparisons between embryos, we set time zero as the frame where we observe the first cellular division post fertilization. Total fluorescence across multiple embryos of the same genotype was averaged and fit to a sigmoidal decay function to determine the maximal fluorescence, the half-life, and the rate of decay (**Fig. 3C-D**). Elevated temperature decreased the total fluorescence in embryos of both genotypes (Fmax_WT_3’UTR-20_ = 6.56 ± 0.08, Fmax_Δ3’UTR-20_ = 4.52 ± 0.06,Fmax_WT_3’UTR-25_ = 4.66 ± 0.04, Fmax_Δ3’UTR-25_ = 3.75 ±0.05). The 3’UTR deletion strain has less maximal fluorescence at time zero than the control strain at both temperatures. Most of the visible GFP::MEX-3 is gone by 60 minutes in both genotypes under all conditions.

However, the half-life of GFP::MEX-3 appears to be shorter for 3’UTR mutant animals at elevated temperature compared control animals (T_1/2_WT_3’UTR-25_ = 35.6±0.4 min, T_1/2_ Δ3’UTR-25_ = 23.7±0.7 min). At standard temperature, the half-lives are similar between genotypes (T_1/2_WT_3’UTR-20_ = 32.9 ± 0.6 min, T_1/2_ Δ3’UTR-20_ = 29.0 ± 0.9 min). The results demonstrate that loss of the 3’UTR reduces the amount of GFP::MEX-3 in embryos and decreases the half-life. They also show that elevated temperature reduces GFP::MEX-3 abundance.

### Morphogenesis fails in the 3’UTR deletion mutant during thermal stress

To understand the nature of the terminal phenotype, we imaged *gfp::3xflag*::*mex-3*::WT 3’UTR and *gfp::3xflag*::*mex-3*::Δ3’UTR mutant embryos as a function of embryonic stage. We compared the results to WT 3’UTR worms cultured on control RNAi or MEX-3 targeting RNAi food. GFP::MEX-3 was observed in young embryos produced from both strains grown on control RNAi food but not *mex-3* RNAi food, demonstrating effective knock down of *mex-3* (**Fig. 4A**). Most embryos produced by worms cultured on control RNAi food appeared normal irrespective of genotype (WT 3’UTR = 96%, Δ3’UTR = 91%). As expected, most embryos produced by worms cultured on *mex-3* RNAi food arrested prior to morphogenesis, consistent with the reported null phenotype (WT 3’UTR = 95% arrested, Δ3’UTR = 93% arrested). Similarly, when *gfp::3xflag*::*mex-3*::Δ3’UTR embryos were grown at the permissive temperature of 20 ºC, embryos appeared normal until after the comma stage when morphogenesis begins (**Fig. 4B-C**). After that, approximately 40% of embryos appeared to arrest, forming terminal embryos with disorganized tissue and zones of apoptosis. The phenotype became more penetrant at elevated temperature, with 100% of late stage embryos appearing abnormal while the younger embryos appeared normal (**Fig. 4B-C**). Interestingly, we noted visual differences between the terminal embryos produced by *mex-3* RNAi compared to the *mex-3* Δ3’UTR deletion mutant, suggesting that the terminal phenotype may be different.

**Figure 4.**
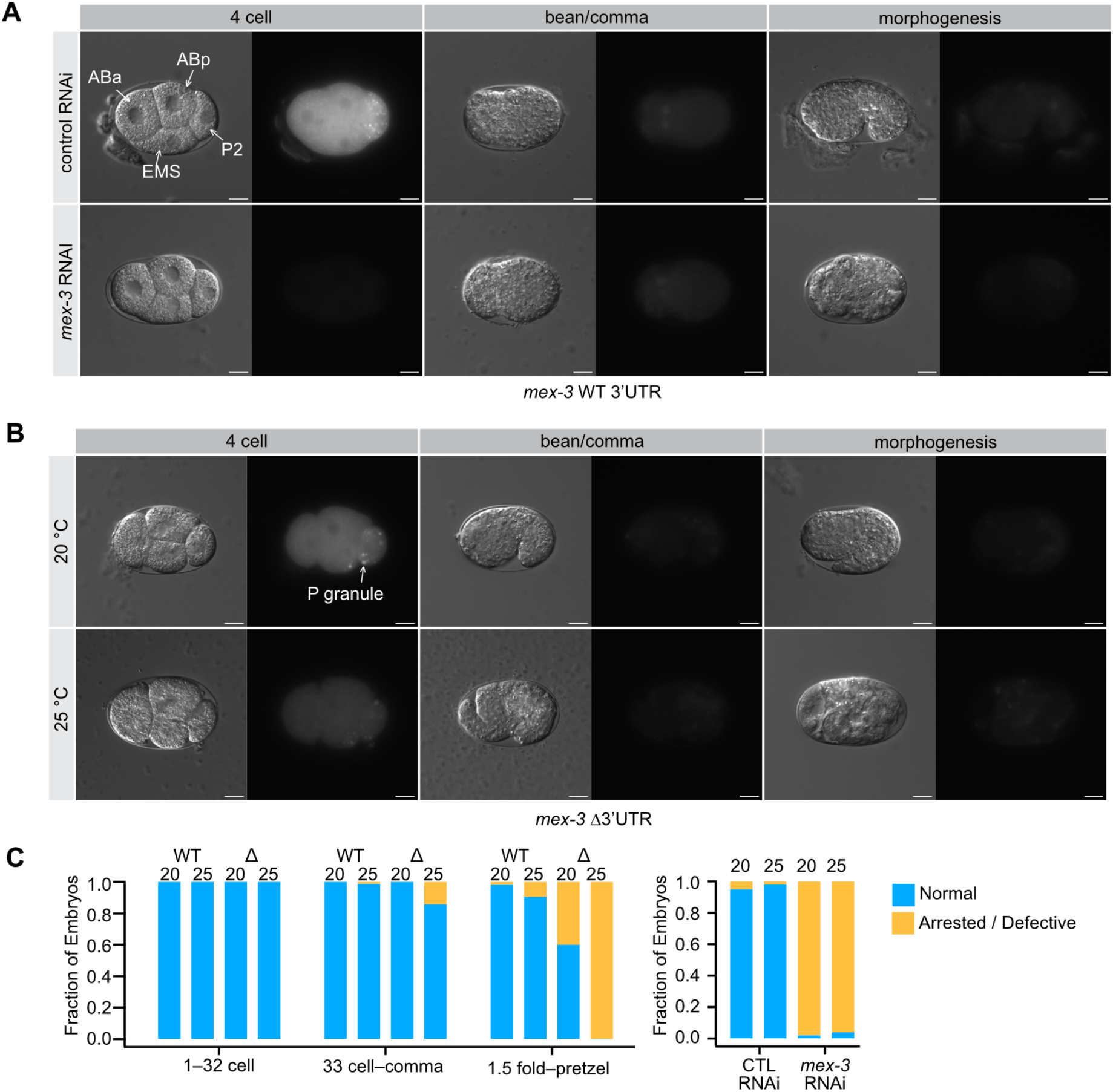
Terminal phenotypes of *mex-3* Δ3’UTR mutants at elevated temperature. **A**. *mex-3* RNAi results in dead embryos that fail at the morphogenesis stage. The phenotype includes expanded muscle lineage cells as well as a few extra germline progenitor cells and disorganized pharyngeal tissue. Scale bars denote 10 microns. **B**. At elevated temperature, the *mex-3* Δ3’UTR mutants appear defective by the comma stage, and terminal embryos that appear different from *mex-3* RNAi. **C**. Fraction of embryos that appear normal vs. abnormal binned by developmental stage. For *mex-3* RNAi animals, arrest happens prior to the comma stage, so all embryos were analyzed together.

To better compare the extent of differentiation in the terminal phenotypes, we crossed the GFP::*mex-3*::Δ3’UTR mutant and the wild type control into OD1854, a germ layer reporter strain. This strain expresses a *Ppha-4*::*pha-4::gfp::pha-4 3’UTR* marker in the endoderm (pharynx and intestine, green), *Phlh-1::gfp::his-72* and *Phlh-1:mCherry::his-72* reporters in mesoderm (body wall muscle, yellow), and *Pdlg1-7I*::*mCherry::his-72* and a *Pcnd-1*::*mCherry::his-72* in the ectoderm (epidermis and about one third of neurons, red, **Fig. 5**) (Wang et al., 2019). In addition to providing a clear view of all three germ layers, the mesoderm reporter specifically marks body wall muscle cells which have been shown to be expanded in *mex-3* null mutants (mex = muscle excess) (Draper et al., 1996).

**Figure 5.**
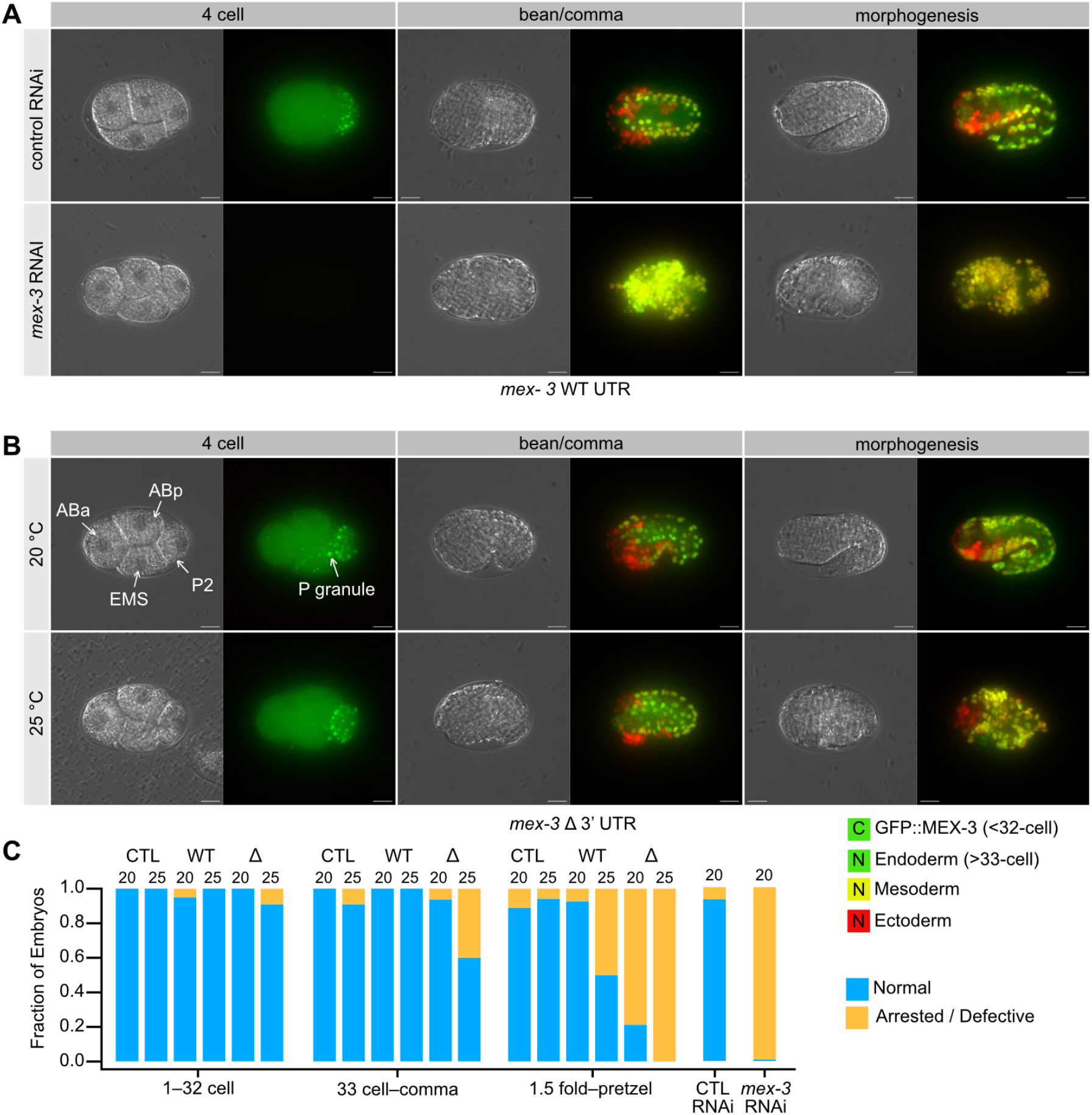
Germ layer patterning defects in the *mex-3* Δ3’UTR mutant. **A**. *mex-3* RNAi results in dead embryos with a large expansion of muscle lineage cells (mesoderm, yellow). Control animals develop normally, with all three germ layers present and appropriately patterned. Scale bars denote 10 microns. **B**. At elevated temperature, the *mex-3* Δ3’UTR mutants form all three germ layers, but patterning appears defective with a moderate expansion of muscle lineage cells (yellow). **C**. Fraction of embryos that appear normal vs. abnormal binned is in figure 4. The genotype and the temperature of growth are shown. CTL is a control for the germ layer background (strain OD1854).

Under standard growth conditions, embryos with the GFP::*mex-3*::WT 3’UTR allele in the germ layer reporter background developed normally. GFP::MEX-3 can be readily distinguished from the transgenic germ layer reporters because 1) its expression is cytoplasmic whereas the reporter transgenes are nuclear, and 2) GFP::MEX-3 disappears prior to transgene expression after zygotic gene activation. Culturing this strain on *mex-3* RNAi food caused a strong and highly penetrant expansion of mesoderm in the terminal embryos (**Fig. 5A**, 100% of embryos). By contrast, 96% of the embryos produced by worms cultured on control RNAi appeared normal with all three germ layers present. Embryos produced by this strain appeared normal under standard growth conditions and at 25 ºC.

In contrast, worms with the *gfp::3xflag*::*mex-3*::WT 3’UTR allele displayed germ layer patterning defects in half of the embryos by the 1.5-fold stage or older at elevated temperature, suggesting the germ layer reporter background causes a synthetic phenotype with GFP::MEX-3. However, the Δ3’UTR mutant showed a much more penetrant germ layer patterning phenotype that manifested earlier and at both temperatures (**Fig. 5B-C**). At 20 ºC, patterning of the Δ3’UTR mutant appeared normal until the 1.5-fold stage wherein 80% of the embryos appeared defective. At 25 ºC, 100% of older Δ3’UTR mutant embryos displayed the patterning phenotype, and 40% of younger embryos between the 32-cell stage and the comma stage also appeared defective. Interestingly, the germ layer patterning phenotype of the Δ3’UTR mutant is different than the *mex-3* RNAi phenotype. All three germ layers are visible, but the pattern is disorganized, and the mesoderm expansion is not as pronounced. As such, the UTR deletion allele potentially retains some MEX-3 activity in the embryo despite reduced abundance and progresses further before arresting compared to *mex-3* RNAi treated animals.

### The number of germline progenitor cells increases in the 3’UTR deletion mutant

A second key feature of the *mex-3* null mutant phenotype is an increase in the number of germline progenitors. In normal animals, two germline progenitors (Z2 and Z3) are produced by the 88-cell stage (Wang and Seydoux, 2013). In *mex-3* null mutants, three or four germline progenitor cells are observed (Draper et al., 1996). To assess whether the Δ3’UTR mutant causes a similar phenotype, we tagged the germline-specific *pgl-1* gene with mCherry at the N-terminus in both the wild type and UTR deletion mutant strains (Kawasaki et al., 1998) (**Fig. 6A**). We note that the *mex-3* WT 3’UTR strain has the expected number of two germline progenitors at 20 or 25 ºC. By contrast, the 3’UTR deletion mutation showed an expanded progenitor cell phenotype in some embryos, but with low penetrance (4% of embryos at 20 ºC., 7% of embryos at 25 ºC, **Fig. 6B**). When the WT 3’UTR strain was cultured on *mex-3* RNAi food, 51% of embryos showed an expansion of germline progenitors, while none of the embryos showed expanded progenitors on control RNAi food. The results show that the 3’UTR mutant can produce the same phenotype as the null, but with low penetrance at both temperatures. This result is consistent with the model that the *mex-3* 3’UTR deletion mutant retains some MEX-3 activity in embryos despite reduced expression.

**Figure 6.**
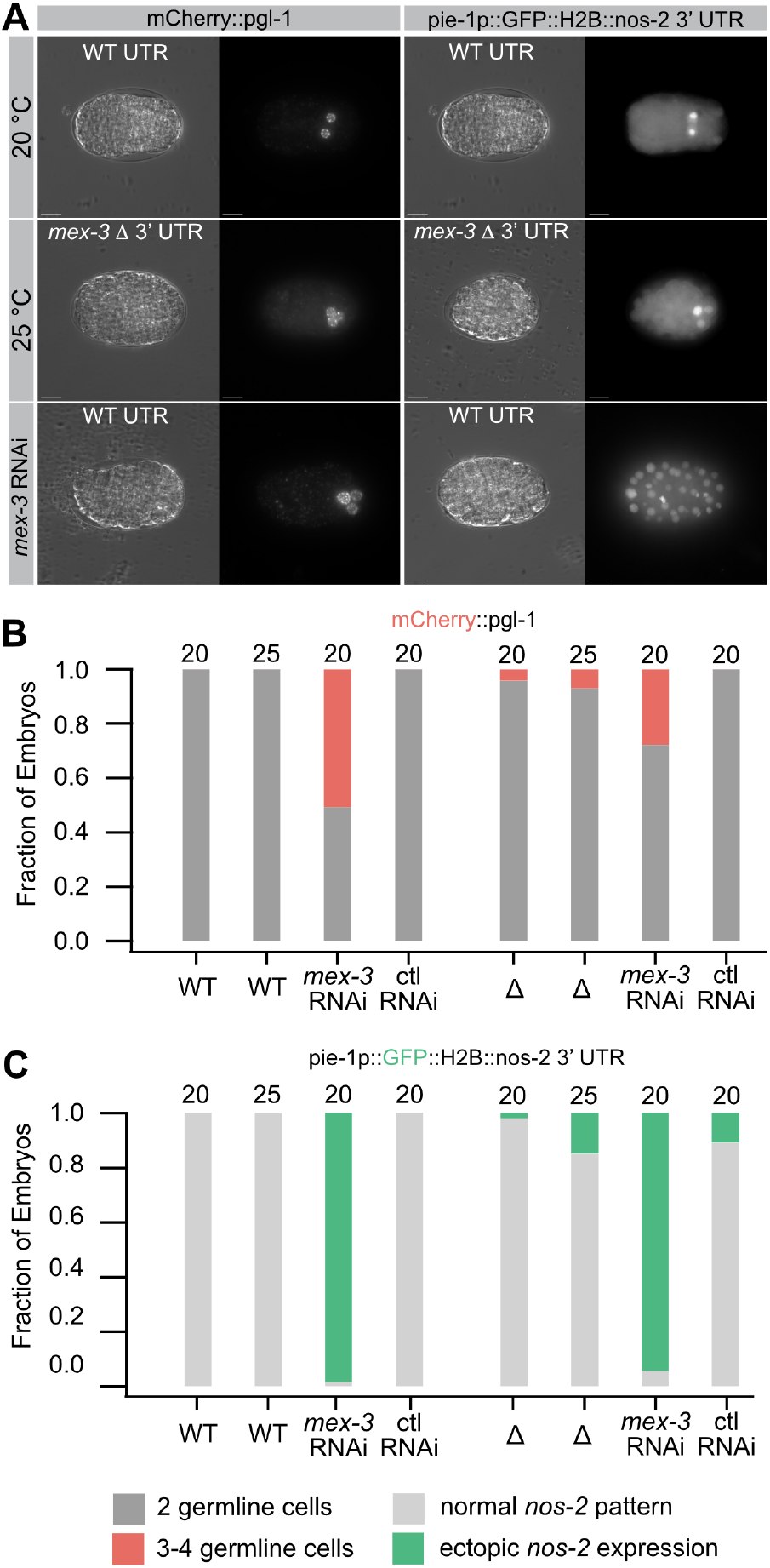
Germline progenitor cell expansion in the *mex-3* 3’UTR deletion. **A**. Two different germline progenitor cell markers, mCherry::*pgl-1* and pie1p::GFP::H2B::*nos-2* 3’UTR, were used to evaluate the number of germline progenitor cells in *mex-3* WT and 3’UTR deletion embryos. MEX-3 directly regulates *nos-2* through its 3’UTR, so expansion to all cells in *mex-3* RNAi is consitent with derepression (D’Agostino et al., 2006). **B**. Quantification of the fraction of embryos displaying 3-4 germline progenitor cells in the mCherry::*pgl-1* background. **C**. Fraction of embryos with >2 germline progenitor cells in pie-1p::GFP::H2B::*nos-2* 3’UTR. reporter.

### The MEX-3 target mRNA nos-2 remains repressed during thermal stress

MEX-3 translationally represses the expression of the germline cell fate determinant *nos-*2 (D’Agostino et al., 2006). We previously showed that MEX-3 regulates a *nos-2* 3’UTR reporter transgene in somatic cells by binding to two MEX-3 Recognition Elements (MREs) in the *nos-2* 3’UTR (Pagano et al., 2009). In that study, *mex-3* RNAi led to ectopic expression of the reporter transgene in all cells of the embryo.

To test whether the *nos-2* reporter remains repressed when the *mex-3* 3’UTR is deleted, we crossed the *nos-2* reporter transgene into the *mex-3* WT 3’UTR strain and the *mex-3* Δ3’UTR strain. We observe ectopic expression limited to 3-4 cells in 2% of embryos produced by the 3’UTR deletion mutant embryos cultured at 20 ºC (**Fig. 6A, C**). This number increases to 15% at 25 ºC. By contrast, we see no ectopic expression in the WT UTR strain at either temperature. We note that expression is not observed in all cells of the early embryo but is instead restricted to 3-4 germline progenitor cells. This contrasts with the *mex-3* RNAi phenotype. Consistent with our previous report, when we culture either strain on *mex-3* RNAi food we observe ectopic expression in all embryonic cells in 95% of *mex-3* WT 3’UTR embryos and 98% of *mex-3* Δ3’UTR embryos (**Fig. 6A, C**), (Pagano et al., 2009). Embryos produced by animals cultured on control RNAi food developed normally. The results suggest that *nos-2* remains repressed in the *mex-3* Δ3’UTR strain in the majority of somatic blastomeres at both temperatures. Though we do see an increase in the number of embryos with limited ectopic expression of the *nos-2* reporter, this could be explained by the increase in the number of germline progenitor cells produced in this mutant, as noted with the *mCherry::pgl-1* marker (**Fig. 6A**). The results reveal another difference between the *mex-3* 3’UTR deletion mutant and the *mex-3* null phenotype, suggesting that the former retains some MEX-3 activity in the embryo despite decreased abundance and the embryonic lethality observed at 25 ºC.

### Dysregulation of mex-3 impacts multiple gene categories

To gain insight into how the 3’UTR deletion mutant impacts gene expression, we used RNA-seq to measure the transcriptome of mutant and control strains at both permissive (20ºC) and restrictive (25ºC) temperatures (**Fig. 7A, B**). We used DeSeq2 to identify differentially expressed genes between the four groups, comparing data for three biological replicates for each strain at each condition (Love et al., 2014).

**Figure 7.**
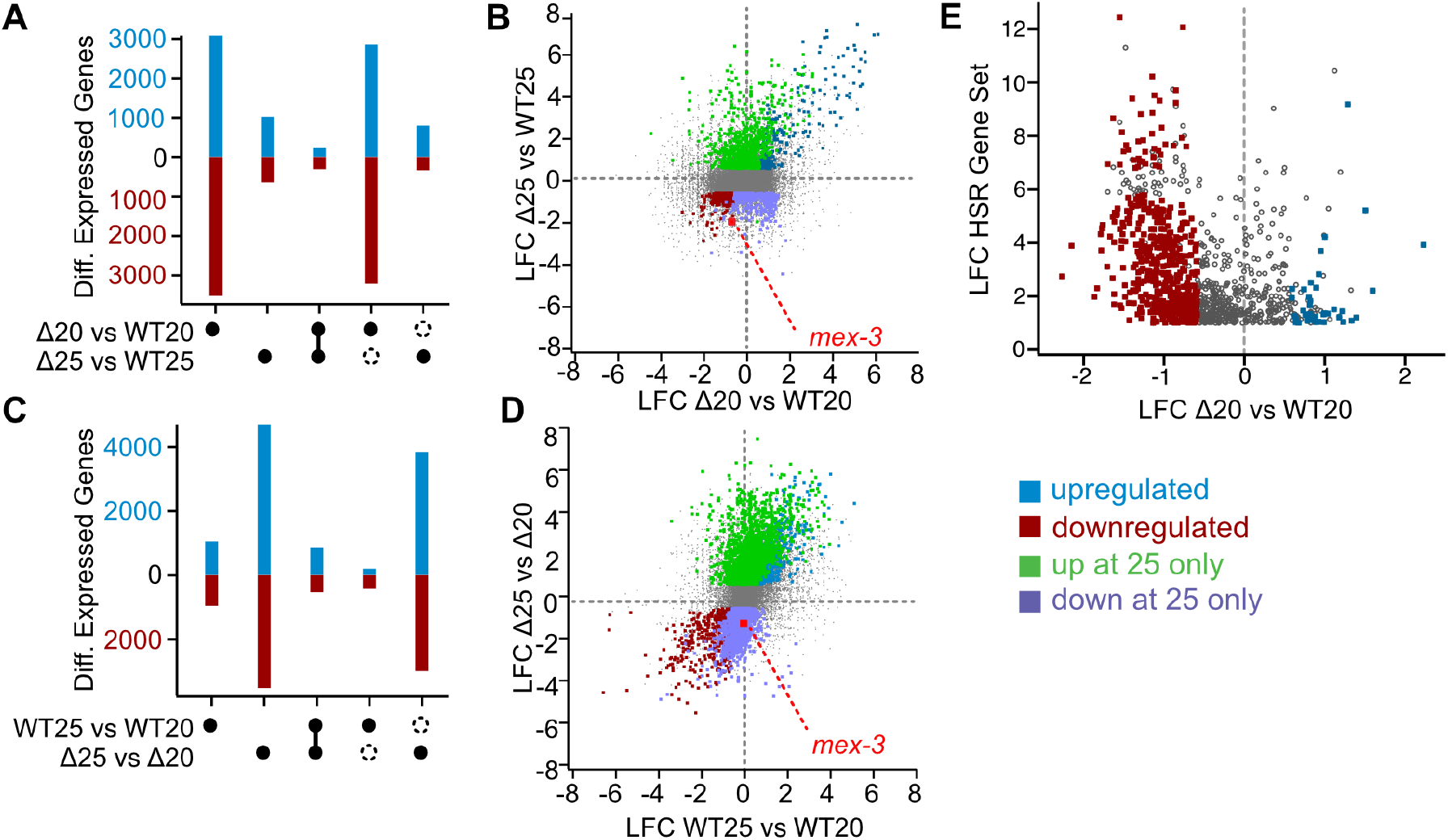
RNA-seq analysis as a function of temperature **A**.UpSet plot comparing genotype responsive changes at both temperatures. The black circles indicate the data sets analyzed. Connected circles are the intersection of the two groups. Open circles represent data present in one group, but absent in the other. Upregulated genes are in blue, downregulated genes are in red. **B**. Comparison of the log_2_fold changes in expression as a function of genotype. The colored circles are defined by the legend **C**. UpSet plot comparing temperature driven changes in each genotype. Labels are as in panel A. **D**. Comparison of the log_2_fold changes in expression as a function of temperature. Labels are as in panel B. The red dot indicates *mex-3* transcripts. **E**. Comparison of the genotype dependent changes in the *mex-3* Δ3’UTR mutant to a high confidence list of heat shock responsive genes. The colors are as defined in the legend. Gray circles represent heat shock responsive genes that are unchanged in the 3’UTR mutant, or are not statistically signficant in our data set (p_adj_< 0.05).

First, we compared the *mex-3* 3’UTR deletion mutant to the control strain cultured at 20ºC. This analysis identifies 3082 upregulated and 3520 downregulated transcripts with a log_2_ fold-change threshold of 0.585 and a P_adj_ value less than or equal to 0.05 (**Fig. 7A**). When the temperature is increased to 25 ºC, the number of affected genes drops to 1025 upregulated and 638 downregulated transcripts, presumably due to the increase in gene expression noise at higher temperature (**Supplementary Data Set 2**). Most of the gene expression changes observed in the mutant at 25 ºC are not observed at 20 ºC, with 78% (795/1025) of upregulated genes and 52% (333/638) of downregulated genes unique to the mutant strain at elevated temperatures (**Fig. 7A,B**). This is not surprising considering that the embryonic lethality phenotype is only prevalent at elevated temperature. We note that one of the downregulated transcripts is *mex-3* itself, suggesting that the reduced GFP::MEX-3 protein expression observed in oocytes and embryos at elevated temperature (**Fig. 3**) correlates with reduced *mex-3* transcript abundance.

### The mex-3 3’UTR mutant limits the heat shock response

Next, to identify temperature responsive differences in the *mex-3* 3’UTR deletion mutant, we reanalyzed the data comparing genotype matched samples grown at 25ºC directly to samples grown at 20ºC (**Fig. 7C**). The data reveal 4707 upregulated and 3518 downregulated transcripts with a log_2_fold-change threshold of 0.585 and a P_adj_ value less than or equal to 0.05. We also identified 1057 upregulated and 954 downregulated transcripts in the control strain when comparing 25ºC to 20ºC samples directly using the same filtering criteria. Somewhat surprisingly, there is little overlap between the temperature responsive transcriptome in each strain (**Fig. 7D**). Only 18% (856/4707) of upregulated transcripts and 15% (530/3518) of downregulated transcripts are observed in both genotypes. As such, the *mex-3* 3’UTR deletion mutant appears to broadly change which transcripts are responsive to elevated temperature, suggesting that the temperature responsive gene expression program is somehow dysregulated in this mutant.

To better understand which gene sets are dysregulated as a function of temperature, we performed a WormCat gene ontology analysis (**Supplemental Fig. 2**) (Holdorf et al., 2020). The analysis reveals low correspondence between the temperature-responsive gene sets between strains. In the control strain, we observe down regulation of metabolism and proteolysis gene sets, consistent with a mild stress response. We also observed downregulation of the extracellular materials gene set and the stress response gene set. Upregulated gene categories were limited to pseudogenes and unassigned genes.

In contrast, the *mex-3* 3’UTR deletion mutant showed changes in numerous gene categories at elevated temperature. The largest and most significant downregulated gene sets in the mutant are cell cycle, mRNA functions, and ribosome. The upregulated sets include neuronal function, signaling, and transmembrane transport. The number and variety of altered gene sets suggests that MEX-3 dysregulation impacts numerous gene pathways. Consistent with this idea, MEX-3 has previously been shown to be important for maintenance of totipotency in the germline, limiting expression of somatic cell fates in the germline, including inhibition of neuronal lineages (Ariz et al., 2009; Ciosk et al., 2006)

Because of the wide variety of changes, we wondered if the global response to temperature stress was altered in the *mex-3* mutant. We compared the changes in gene expression observed in the *mex-3* 3’UTR deletion mutant to the canonical set of genes upregulated in response to heat shock. A high confidence set of 1060 protein coding genes that are upregulated in wildtype (N2) worms when treated with a heat shock (35 ºC for four hours) has been published (Schreiner et al., 2019). We observe that 546 (52%) of these genes are downregulated in the *mex-3* 3’UTR deletion mutant compared to the genotype matched control worms when grown at standard temperature (**Fig. 7E**). By contrast, only 42 (4%) are upregulated. As such, approximately half of the heat shock responsive genes are expressed at a lower basal level in the *mex-3* 3’UTR deletion mutant than in control worms. The data suggests that MEX-3 helps to establish the baseline heat shock response, and that reduced expression of these genes may contribute to the embryonic lethality observed upon mild temperature increase.

## DISCUSSION

In this study we show that a *mex-3* 3’UTR deletion mutant has a strong and highly penetrant embryonic lethal phenotype when cultured at a temperature of 25 ºC. This phenotype is characterized by reduced GFP::MEX-3 levels in maturing oocytes and embryos, failure to undergo morphogenesis, and an expanded number of embryonic germline progenitor cells. Unlike *mex-3* RNAi embryos, the 3’UTR deletion mutant embryos cultured at elevated temperature generate all three germ layers and show less pronounced expansion of body wall muscle. Moreover, a germline specific *nos-2* 3’UTR reporter remains repressed in somatic blastomeres in the embryos. The results suggest that the 3’UTR deletion mutant is likely a temperature sensitive hypomorphic allele, with reduced MEX-3 activity in embryos that remains sufficient to promote differentiation beyond what is observed in a *mex-3* null allele, but insufficient to support viability.

This result is surprising because the most obvious phenotype of the *mex-3* 3’UTR deletion under standard growth conditions is increased GFP::MEX-3 expression throughout the germline (Albarqi and Ryder, 2021). This suggests two things. First, reduced MEX-3 abundance is worse for reproductive fecundity than elevated expression. Second, a major role of the *mex-3* 3’UTR is to ensure robust MEX-3 expression at the oocyte-to-embryo transition in variable environmental conditions. Though animals can thermoregulate by moving to a more suitable environment, embryos cannot, so it is reasonable to hypothesize that the worm has evolved a mechanism to ensure that a key cell fate determinant is expressed under a variety of environmental conditions.

### Repressing and activating circuits in the mex-3 3’UTR

The results presented here suggest that the *mex-3* 3’UTR contains at least two regulatory circuits that act in different regions of the germline (**Fig. 8A)**. The first is a repressive circuit that ensures *mex-3* silencing in the meiotic region of the germline before recellularization into oocytes. The second is an activating circuit that functions in maturing oocytes and early embryos that boosts MEX-3 levels at the oocyte-to-embryo transition. The repressive circuit does not appear to be temperature sensitive, as de-repression is observed in all conditions in the *mex-3* 3’UTR mutant animals. Failure of the activating circuit appears to be exacerbated by elevated temperature, as mutant animals cultured at 25 ºC have less GFP::MEX-3 in the most proximal oocyte and in embryos than wild-type controls or mutants cultured at standard temperature.

**Figure 8.**
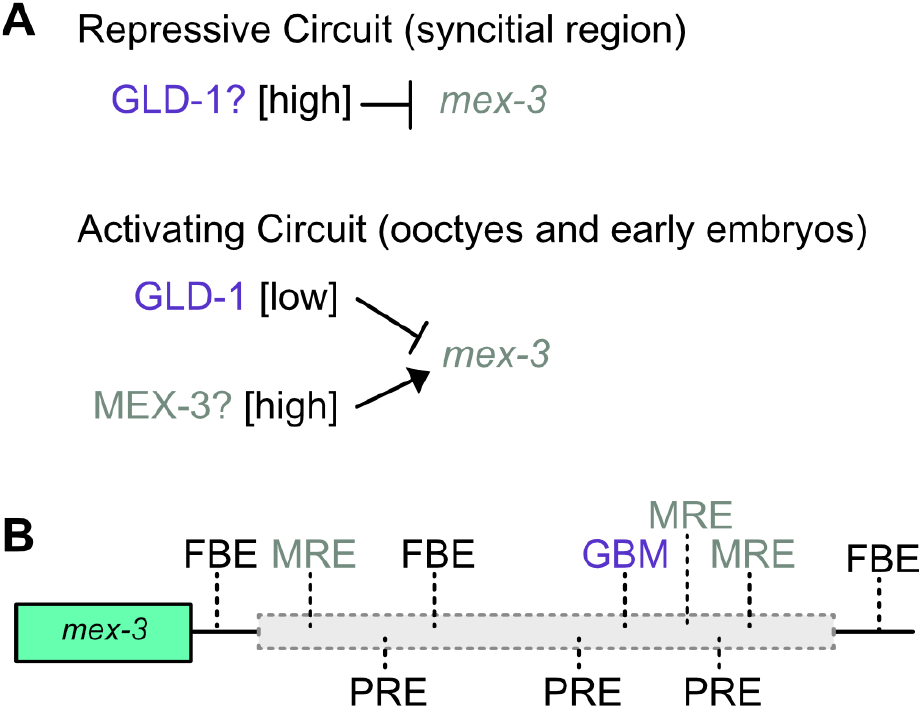
A hypothetical model of repressing and activating circuits encoded by the *mex-3* 3’UTR. **A**. Competition model for how GLD-1 and MEX-3 could contribute to regulation in different regions of the germline. **B**. Distribution of FBF binding elements (FBE), MEX-3 recognition elements (MRE), POS-1 recognition elements (PRE), and GLD-1 binding motifs (GBM) in the *mex-3* 3’UTR.

The trans factors that govern *mex-3* 3’UTR-dependent repression and/or activation are not known, but a few candidates arise from inspection of known RBP motifs in the *mex-3* 3’UTR (**Fig. 8B)**. GLD-1 is a STAR/KH domain RNA-binding protein that binds with high affinity and specificity to a heptanucleotide motif in its target RNAs (Jungkamp et al., 2011; Ryder et al., 2004; Wright et al., 2011). Its expression increases at the switch from mitosis to meiosis, then decreases during oogenesis (Jones et al., 1996). There is a strong GLD-1 binding motif (GBM) in the *mex-3* 3’UTR (Ryder et al., 2004; Wright et al., 2011). This interaction has been confirmed by high throughput GLD-1 interactome studies (Jungkamp et al., 2011; Wright et al., 2011). The anticorrelated pattern of expression, the presence of a high affinity consensus binding motif, and the known role of GLD-1 in translationally repressing other germline transcripts make it a strong candidate for a trans-acting factor that represses *mex-3* in the meiotic germline. However, a 134 base pair deletion that removes the high affinity GBM has no phenotype and does not display altered expression (Albarqi and Ryder, 2021). As such, additional motifs, potentially recognized by other factors, may also be required or may be redundant with the GBM.

The trans acting factors that govern activation in oocytes and embryos are harder to predict. In addition to a strong GBM, the *mex-3* 3’UTR harbors three consensus binding motifs for POS-1 (PREs), three for FBF-1 (FBEs), and three motifs for MEX-3 itself (MREs) (Bernstein et al., 2005; Farley et al., 2008; Kalchhauser et al., 2011; Pagano et al., 2009; Wang et al., 2009). The 3’UTR deletion allele removes all three *mex-3* binding sites, all three POS-1 sites, and one of the FBF sites. POS-1 is a tandem zinc finger RBP expressed exclusively in early embryos after fertilization (Tabara et al., 1999). As such, this protein can’t be responsible for *mex-3* activation in late-stage oocytes. Similarly, FBF is an RBP that is expressed in mitotic progenitor germline progenitor cells where it promotes progenitor fate renewal (Zhang et al., 1997). It is not found in late-stage oocytes and likely doesn’t contribute to enhanced expression.

The remaining candidate is MEX-3 itself. It is intriguing to consider the possibility that MEX-3 might activate its own expression. The three MEX-3 binding sites are in distant regions of the 3’UTR, which could explain why most of the 3’UTR must be removed to observe the phenotype. MEX-3 abundance increases as GLD-1 decreases in oocytes. The GBM is adjacent to two of the MREs, suggesting the potential for competition between MEX-3 and GLD-1 for binding to adjacent binding motifs. It remains equally likely that other RBPs, binding to yet to be determined cis-regulatory motifs, govern increased GFP::MEX-3 abundance in oocytes. More work will be needed to identify the functional elements within the 3’UTR responsible for the phenotypes presented here.

### What is the basis of temperature sensitivity?

The molecular mechanism that governs the temperature sensitivity of the *mex-3* 3’UTR deletion phenotype is not known. Temperature sensitive mutations are often found within the protein coding sequence of genes where the mutation alters protein folding stability (Sandberg et al., 1995). Here, the mutation is present in the 3’UTR and does not perturb the protein coding sequence. As such, the temperature sensitivity must arise by a different mechanism.

The RNA-seq data reveal a few plausible mechanisms. First, a large fraction of the high confidence heat shock response genes is downregulated in the *mex-3* 3’UTR mutant. It could be that dysregulation of *mex-3* activity leads to attenuation of the heat shock response by a direct or indirect mechanism. Loss of the ability to adapt to heat could potentially explain the temperature sensitive phenotype. HSF-1 is a conserved transcriptional regulator of the heat shock response (Hajdu-Cronin et al., 2004). In *C. elegans*, monomeric HSF-1 is held in an inactive state by forming a complex with DDL-1/2 and HSB-1 (Chiang et al., 2012). Upon experiencing heat stress, HSF-1 is released from this complex and forms a homotrimer that binds to heat shock elements found in the promoters of many heat response genes Intriguingly, we note that *ddl-1* and *ddl-2* transcripts are both upregulated in the *mex-3* 3’UTR mutant, suggesting that perhaps MEX-3 controls the abundance of these negative regulators of the heat shock response through some alternative mechanism, perhaps through direct regulation of their mRNA stability, or control of some upstream component that feeds into the HSF pathway. More work will be needed to determine if these factors or others contribute to the embryonic lethality phenotype.

Second, our data show that many ribosomal protein gene transcripts are downregulated in the *mex-3* 3’UTR mutant at elevated temperature. This suggests that increasing temperature is leading to a coordinated decrease in ribosome biogenesis, which could broadly impact maternal gene translation in the embryo including MEX-3 itself, as observed in **Fig. 3**. Ribosome biogenesis is essential to successful reproduction, and maternally supplied ribosomes are sufficient for successful embryogenesis (Cenik et al., 2019). Ribosome protein gene biosynthesis is also necessary for mitochondrial ribosome function and mitochondrial health. A more recent study demonstrated that haploinsufficiency of five different ribosomal protein genes alters mitochondrial morphology, and haploinsufficiency of *rps-10* had broad effects on mitochondrial function and cellular energy states (Surya et al., 2024). We note that ribosomal protein genes and metabolism genes were enriched in our WormCat ontology analysis in the mutant at elevated temperature. How MEX-3 impacts ribosome biogenesis is not known.

Many additional models consistent with these data could be drawn. While it is tempting to speculate that specific gene expression changes that manifest at 25 deg. C in the mutant strain are responsible for the temperature sensitive phenotype, it remains equally likely that these changes are a consequence, rather than a cause of the phenotype. Considerably more work will be necessary to dissect the molecular basis of the temperature sensitive phenotype.

### Temperature sensitivity, developmental processes, and post-transcriptional regulation

Post-transcriptional regulatory pathways have been shown to be critical to biological robustness in other aspects of nematode development as well (Burke et al., 2015; Ilbay and Ambros, 2019). Specifically, microRNAs (miRNAs) that repress key heterochronic pathway mRNA targets during larval development rarely have a phenotype when mutated (Alvarez-Saavedra and Horvitz, 2010; Miska et al., 2007). This is in part due to the redundancy built into microRNA families—multiple miRNAs in the same family recognize the same seed sequence and can thus act redundantly. Also, multiple miRNAs can regulate the same mRNA through distinct, yet functionally redundant, binding sites. As such, their importance could have been easily overlooked. The Abbot lab identified strong miRNA mutant phenotypes in sensitized genetic backgrounds (Brenner et al., 2010). Expanding upon this study, the Ambros lab demonstrated that multiple miRNA family mutants display enhanced phenotypes upon oscillating temperature stress but not upon other forms of stress (Burke et al., 2015; Ilbay and Ambros, 2019).

The results presented here show that the *mex-3* 3’UTR is essential to embryonic morphogenesis at elevated temperature. It is appealing to speculate that a major role for the masked “load” of maternal mRNAs first described by Spirin in 1966 (Spirin, 1966) is to act as a buffer to ensure sufficient maternal protein is produced in uncertain environments. Whether the results presented here are generalizable to other maternal transcripts remains to be seen. Few large 3’UTR deletion alleles in maternal transcripts have been engineered to date, so a comprehensive assessment cannot be made at this time.

## MATERIALS AND METHODS

### Strains and Nematode Culture

All strains used in this study are listed in **Table 1**. Strains were maintained by growing animals on NGM (Nematode Growth Medium) seeded with *E. coli* (OP50) under standard conditions (Stiernagle, 2006). All primers used to produce or evaluate the strains are listed in **Supplemental Table 1**. WRM75 was generated by crossing WRM52 males with OD1854 (germ layer reporter) hermaphrodites, selecting heterozygous male cross progeny, then back crossing the F1 males to strain OD1854 before isolating individuals and following progeny, confirming the desired genotype by PCR and fluorescence imaging. WRM77 was made by the same approach using DG4269 males.

**Table 1:**
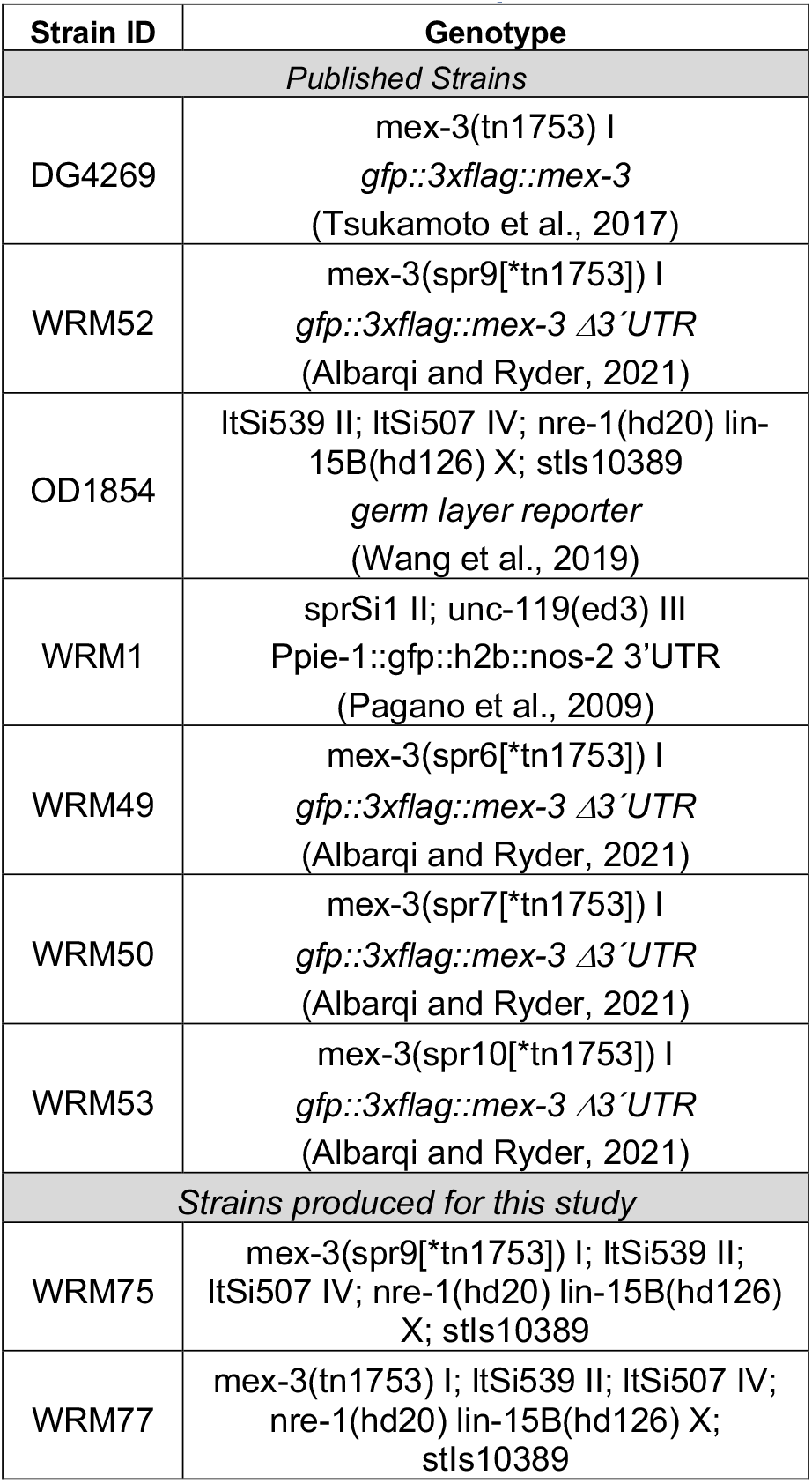
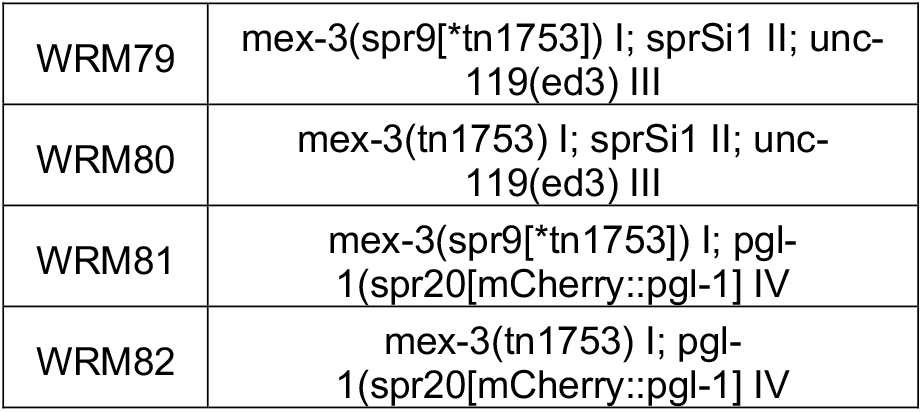
Strains used in this report.

| Strain ID | Genotype |
| --- | --- |
| *Published Strains* |  |
| DG4269 | mex-3(tn1753) I<br>*gfp::3xflag::mex-3*<br>(Tsukamoto et al., 2017) |
| WRM52 | mex-3(spr9[*tn1753]) I<br>*gfp::3xflag::mex-3 Δ3'UTR*<br>(Albarqi and Ryder, 2021) |
| OD1854 | ItSi539 II; ItSi507 IV; nre-1(hd20) lin-15B(hd126) X; stIs10389<br>*germ layer reporter*<br>(Wang et al., 2019) |
| WRM1 | sprSi1 II; unc-119(ed3) III<br>Ppie-1::gfp::h2b::nos-2 3'UTR<br>(Pagano et al., 2009) |
| WRM49 | mex-3(spr6[*tn1753]) I<br>*gfp::3xflag::mex-3 Δ3'UTR*<br>(Albarqi and Ryder, 2021) |
| WRM50 | mex-3(spr7[*tn1753]) I<br>*gfp::3xflag::mex-3 Δ3'UTR*<br>(Albarqi and Ryder, 2021) |
| WRM53 | mex-3(spr10[*tn1753]) I<br>*gfp::3xflag::mex-3 Δ3'UTR*<br>(Albarqi and Ryder, 2021) |
| *Strains produced for this study* |  |
| WRM75 | mex-3(spr9[*tn1753]) I; ItSi539 II;<br>ItSi507 IV; nre-1(hd20) lin-15B(hd126) X; stIs10389 |
| WRM77 | mex-3(tn1753) I; ItSi539 II; ItSi507 IV;<br>nre-1(hd20) lin-15B(hd126) X;<br>stIs10389 |
| WRM79 | mex-3(spr9[*tn1753]) I; sprSi1 II; unc-119(ed3) III |
| WRM80 | mex-3(tn1753) I; sprSi1 II; unc-119(ed3) III |
| WRM81 | mex-3(spr9[*tn1753]) I; pgl-1(spr20[mCherry::pgl-1] IV |
| WRM82 | mex-3(tn1753) I; pgl-1(spr20[mCherry::pgl-1] IV |

WRM79 was generated by crossing WRM52 males with WRM1 (*nos-2* 3’UTR reporter) hermaphrodites, then backcrossing the F1 males to strain WRM1 before isolating individuals and following progeny, confirming the desired genotype by PCR. WRM80 was made by the same approach using DG4269 males.

WRM81 and WRM82 (*pgl-1* reporter strains) were generated using CRISPR/Cas9 following the procedure of Ghanta and Mello (Ghanta et al., 2021; Ghanta and Mello, 2020). To produce WRM81, WRM52 worms were injected with an RNP mixture containing *Streptococcus pyogenes* Cas9 (IDT #1081058) (10µg/µL), tracrRNA (0.4 µg/µL), crRNA (0.4 µg/µL; gRNA sequence: 5’-GUUUCAUCCAUUUCACAUGG -3’), 500 ng of melted dsDNA recombination template (25 ng/µL final conc.), PRF::rol-6 plasmid (500 ng/µL) and nuclease free water. The recombination template was made by PCR amplifying the recombination template from a plasmid template (pCCM953, a gift from Craig Mello). The PCR product was purified using a kit (Zymo Research DNA Clean & Concentrator™ Cat. No. D4034) The desired genotype was confirmed by PCR and sequencing. WRM82 was made by injecting DG4269 animals with the same injection mix.

### Brood Size and Hatch Rate Assays

Brood size and hatch rate measurements were conducted in triplicate for each condition and genotype. Strains were synchronized by bleaching adult worms in a 20% alkaline hypochlorite solution (3 mL concentrated Clorox bleach, 3.75 mL 1M filtered sodium hydroxide, 8.25 mL filtered MilliQ H2O), embryos were recovered by centrifugation, then washed extensively in M9 buffer before they were allowed to hatch overnight. Once synchronized as starved L1 larvae, animals were transferred onto NGM agar plates seeded with OP50 and allowed to grow to the L3/L4 stage at room temperature. For each replicate, 20 individual L4 animals were separated onto individual NGM plates then introduced to a stress condition (15°C, 25 °C, 300 mM NaCl, *P. aeruginosa* PA14) or a standard growth condition control (20 ºC, 50 mM NaCl, *E. coli* OP50). After 24 hours, each animal was transferred to a fresh plate, the total number of embryos laid was counted, then both plates were reintroduced to the stress condition. After another 24 hours, the number of hatched larvae were also counted from the first plate. The process was repeated until the animals no longer produced fertilized embryos (∼ 7 days post synchronization). To measure recovery from 25 ºC stress, assays were performed as above but worms were placed in the stress condition for 24 hours, then removed to 20 ºC for the remainder of the experiment.

From these data, the total number of embryos produced, the total viable progeny produced, and the hatch rate (fraction of viable embryos) were determined for each genotype, condition, or recovery period. Statistical significance of differences was assessed by one way ANOVA with Bonferroni correction for multiple hypothesis testing using StatPlus software (AnalystSoft, Brandon, FL).

For the pathogenic stress assays, NGM plates were prepared with increased peptone (3.5 g/L, standard is 2.5 g/L). Plates were seeded with 15 μL of a 1:5 dilution of a saturated overnight *P. aeruginosa* PA14 culture. Seeded plates were incubated at 37 °C for 24 hours, moved to 25 °C for another 24 hours, then stored at 4 °C for a maximum of a week before use.

### RNA interference

RNAi by feeding was performed by feeding as previously described (Conte and Mello, 2003). RNAi plates were prepared by supplementing NGM agar with ampicillin and Isopropyl β-D-1-thiogalactopyranoside (IPTG; 100mM Ampicillin and 1mM IPTG) before pouring. RNAi plates were seeded with *E. coli* HT115 transformed with plasmid that expresses dsRNA targeting *mex-3* or with an empty vector control as previously described (Pagano et al., 2009). Animals were synchronized as L1 larvae, seeded onto RNAi plates, and allowed a minimum of 24 hours to feed on MEX-3 RNAi food before imaging or scoring for phenotypes. The effectiveness of the RNAi targeting *mex-3* was confirmed by loss of *gfp::3xflag::mex-3* expression in oocytes and embryos, and by scoring for the strong embryonic lethality phenotype.

### Embryo Imaging and Phenotyping

Embryos from DG4269, WRM52, WRM75, or WRM77 were recovered from young adults by dissection. Older and terminal embryos were recovered from plates by washing with M9, then repeated centrifugations (3000 rpm) and washes with M9 to isolate embryos from bacteria. Embryos were mounted on 2% agarose pads on glass slides and kept hydrated under the cover slip with 3-5 μL of M9 solution. Images of individual embryos were collected in twenty layers along the Z axis using a Zeiss AxioObserver 7 microscope with differential interference contrast and fluorescence imaging optics, a 63X objective, and a Zeiss Axiocam 506 mono camera. Fiji software (ImageJ v. 2.9.0) was used to create a Z projection for each fluorescence channel. For strains expressing multiple fluorescent proteins, all fluorescence channels were merged into a single image for analysis. The genotype and growth condition were documented by one lab member, and the state of the embryo (normal, abnormal, or dead) was scored blind by another. The fraction of normal embryos was determined by calculating the number of apparently normal embryos divided by the total number of living embryos.

Young embryos from WRM79, WRM80, WRM81, and WRM82 were recovered and imaged as above. Only embryos older than the 88-cell stage were considered. The total number of cells expressing *Ppie-1::gfp::h2b::nos-2-3’UTR* or *mCherry::pgl-1* were counted. The fraction of normal embryos was determined by calculating the number of embryos with expression in two cells (normal) divided by the total number of embryos scored.

### Embryogenesis Movies

Embryos were recovered from young adult hermaphrodites and mounted on 2% agarose pads for imaging as described above. Only embryos that were younger than the 2-cell stage were imaged. DIC and GFP images were collected every five to seven minutes. Focus was maintained throughout imaging using a Zeiss Definite Focus unit. Movies were analyzed using Fiji / ImageJ software first by cropping the 16-bit image stacks to 40 × 60 microns centering the embryo, then measuring the mean GFP::MEX-3 fluorescence across the entire embryo per image in the stack using the “Plot Z-axis Profile” tool. Because embryogenesis cannot be synchronized, time zero was arbitrarily defined as the first frame that showed completed cytokinesis of the first cell division, enabling comparison between multiple embryos. The mean fluorescence intensity was normalized by dividing by the local minimum intensity across all frames of the stack. The average mean fluorescence per time point across at least three embryos was plotted, and the data fit to the following sigmoidal decay equation using IgorPro 9.0.2 (Oswego, OR):

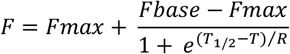

where F is the mean GFP intensity across all frames, Fmax is the maximal intensity, Fbase in the minimum intensity, T is the time (in minutes), T_1/2_ is the half-life, and R is the rate of decay. The standard deviation of the fitted parameters was determined from a global fit of all data per genotype and condition.

### Young adult imaging and quantitation

Young adult animals were picked onto a 2% agarose pad on a glass slide in M9 supplemented with levamisole (1 mM) to immobilize them. DIC and GFP images of the germline of intact worms were collected with a Zeiss AxioObserver 7 microscope with a 20X objective and a Zeiss Axiocam 506 mono camera. GFP fluorescence in oocytes was calculated by drawing a line with 30 pixel width across each of the four most proximal oocytes using Fiji software, avoiding the nucleus. The mean fluorescence intensity within the line was measured and the background subtracted for each oocyte. For each genotype and condition, the background corrected fluorescence intensity for each of the four most proximal oocytes was calculated across at least twelve animals. Differences between the distribution of intensities between oocytes, genotypes, and conditions were compared using a one-way ANOVA with Bonferroni correction for multiple hypothesis testing to assess statistical significance.

### RNA sequencing

DG4269 and WRM52 animals were cultured at 20°C and 25°C and harvested from unstarved plates with filtered nuclease-free water. Each population was washed at least 3 times with water prior to RNA isolation using trizol, chloroform, and isopropanol. Ribosomal RNA from each sample was depleted using a rRNA depletion protocol for *C. elegans* (Duan et al., 2020). Library prep for RNA-seq was performed using the NEBNext Ultra II library prep kit (cat #E7775S) following the manufacturer’s protocol. NEBNext Multiplex Oligos for Illumina (Dual Index Primer Set 1) (cat #E7600) was used for library indexing. Concentrations for each library was determined using both Qubit and fragment bioanalyzer. Barcoded libraries were sequenced using an Illumina NEXTSeq 1000.

Raw sequencing reads were mapped to the *C. elegans* genome build WBcel135 using the OneStopRNAseq pipeline (Li et al., 2020). Briefly, FastQC v0.11.5 and MultiQC v1.6 were used for raw and post-alignment quality control, respectively (Andrews, 2010; Ewels et al., 2016). STAR v2.7.5a was used to align the reads to the reference genome using WBcel235.90 annotations with default settings except for the following paramaters ‘-Q 20 –minOverlap 1 --fracOverlap 0 -p -B -C’ for paired-end strict-mode analysis (Dobin et al., 2013). Differential expression (DE) analysis was performed with DESeq2 v1.28.1 (Love et al., 2014). Significantly differentially expressed genes were filtered with the criteria FDR < 0.05 and absolute log2 fold change (|LFC|) > 0.585. The data are available in **Supplementary Data Set 2**.

## Supporting information

Supplemental Data Set 1

Supplemental Data Set 2

## Data availability statement

All of the data and statistical analyses presented in in figure 2 through figure 7 of this work are available in **Supplemental Data Sets 1** and **2**. All sequencing data have been uploaded to the NCBI Sequence Read Archive under Bioproject PRJNA1185698.

## ACKNOWLEDGEMENTS

The authors thank Dr. Krishna Ghanta and Dr. Ebru Kaymak for technical support and helpful discussions concerning strain generation and characterization. We thank Dr. Takao Ishidate and Dr. Craig Mello for the gift of plasmid pCCM953 used to make the *mCherry::pgl-1* strains. We thank Dr. Ye Duan for technical support with the rRNA depletion protocol. We thank Qiangzong Yin and Dr. Pablo Bora for technical assistance with RNA sequencing. We thank Dr. Zeynep Mirza and Dr. Victor Ambros for helpful discussions and reagents for the experiments with *Pseudomonas aeruginosa* PA14. We thank Dr. Rebecca Green for the gift of nematode strain OD1854. We thank Sharon Noronha for helpful comments on this manuscript. This work was supported by NIH Grants R01GM145062 and R01HD111505 to S.P.R.

## AUTHOR CONTRIBUTIONS

HB and SR conceived, designed, and interpreted the experiments in this work. HB performed all experiments except for the 15° C, osmotic stress, and *Pseudomonas aeruginosa* brood size assays, which were performed by both HB and HV. The GFP::MEX-3 oocyte measurements which were performed by both SK and HB. HB and SR wrote the manuscript in consultation with all authors.

## REFERENCES

Abram, P. K., Boivin, G., Moiroux, J. and Brodeur, J. (2017). Behavioural effects of temperature on ectothermic animals: unifying thermal physiology and behavioural plasticity. Biol Rev Camb Philos Soc 92, 1859–1876.

Albarqi, M. M. Y. and Ryder, S. P. (2021). The endogenous mex-3 3 UTR is required for germline repression and contributes to optimal fecundity in C. elegans. PLoS Genet 17, e1009775.

Albarqi, M. M. Y. and Ryder, S. P. (2022). The role of RNA-binding proteins in orchestrating germline development in Caenorhabditis elegans. Front Cell Dev Biol 10, 1094295.

Alvarez-Saavedra, E. and Horvitz, H. R. (2010). Many families of C. elegans microRNAs are not essential for development or viability. Curr Biol 20, 367–373.

Anderson, J. L., Albergotti, L., Ellebracht, B., Huey, R. B. and Phillips, P. C. (2011). Does thermoregulatory behavior maximize reproductive fitness of natural isolates of Caenorhabditis elegans? BMC Evolutionary Biology 11, 157.

Andrews, S. (2010). FastQC: A quality control tool for high throughput sequence data. https://www.bioinformatics.babraham.ac.uk/projects/fastqc/.

Aprison, E. Z. and Ruvinsky, I. (2014). Balanced Trade-Offs between Alternative Strategies Shape the Response of C. elegans Reproduction to Chronic Heat Stress. PLOS ONE 9, e105513.

Ariz, M., Mainpal, R. and Subramaniam, K. (2009). elegans RNA-binding proteins PUF-8 and MEX-3 function redundantly to promote germline stem cell mitosis. Dev Biol 326, 295–304.

Begasse, M. L., Leaver, M., Vazquez, F., Grill, S. W. and Hyman, A. A. (2015). Temperature Dependence of Cell Division Timing Accounts for a Shift in the Thermal Limits of C. elegans and C. briggsae. Cell reports 10, 647–653.

Bernstein, D., Hook, B., Hajarnavis, A., Opperman, L. and Wickens, M. (2005). Binding specificity and mRNA targets of a C. elegans PUF protein, FBF-1. RNA 11, 447–458.

Brenner, J. L., Jasiewicz, K. L., Fahley, A. F., Kemp, B. J. and Abbott, A. L. (2010). Loss of individual microRNAs causes mutant phenotypes in sensitized genetic backgrounds in C. elegans. Curr Biol 20, 1321–1325.

Bull, J. J. and Vogt, R. C. (1979). Temperature-dependent sex determination in turtles. Science 206, 1186–1188.

Burke, S. L., Hammell, M. and Ambros, V. (2015). Robust Distal Tip Cell Pathfinding in the Face of Temperature Stress Is Ensured by Two Conserved microRNAS in Caenorhabditis elegans. Genetics 200, 1201–1218.

Byerly, L., Cassada, R. C. and Russell, R. L. (1976). The life cycle of the nematode Caenorhabditis elegans. I. Wild-type growth and reproduction. Dev Biol 51, 23–33.

Cenik, E. S., Meng, X., Tang, N. H., Hall, R. N., Arribere, J. A., Cenik, C., Jin, Y. and Fire, A. (2019). Maternal Ribosomes Are Sufficient for Tissue Diversification during Embryonic Development in C. elegans. Dev Cell 48, 811–826 e816.

Chiang, W. C., Ching, T. T., Lee, H. C., Mousigian, C. and Hsu, A. L. (2012). HSF-1 regulators DDL-1/2 link insulin-like signaling to heat-shock responses and modulation of longevity. Cell 148, 322–334.

Ciosk, R., DePalma, M. and Priess, J. R. (2004). ATX-2, the C. elegans ortholog of ataxin 2, functions in translational regulation in the germline. Development 131, 4831–4841.

Ciosk, R., DePalma, M. and Priess, J. R. (2006). Translational regulators maintain totipotency in the Caenorhabditis elegans germline. Science 311, 851–853.

Conover, D. O. and Heins, S. W. (1987). Adaptive variation in environmental and genetic sex determination in a fish. Nature 326, 496–498.

Conte, D., Jr. and Mello, C. C. (2003). RNA interference in Caenorhabditis elegans. Current protocols in molecular biology / edited by Frederick M. Ausubel … [et al.] Chapter 26, Unit 26 23.

D’Agostino, I., Merritt, C., Chen, P. L., Seydoux, G. and Subramaniam, K. (2006). Translational repression restricts expression of the C. elegans Nanos homolog NOS-2 to the embryonic germline. Dev Biol 292, 244–252.

Dobin, A., Davis, C. A., Schlesinger, F., Drenkow, J., Zaleski, C., Jha, S., Batut, P., Chaisson, M. and Gingeras, T. R. (2013). STAR: ultrafast universal RNA-seq aligner. Bioinformatics 29, 15–21.

Draper, B. W., Mello, C. C., Bowerman, B., Hardin, J. and Priess, J. R. (1996). MEX-3 is a KH domain protein that regulates blastomere identity in early C-elegans embryos. Cell 87, 205–216.

Dreier, J. W., Andersen, A. M. and Berg-Beckhoff, G. (2014). Systematic review and meta-analyses: fever in pregnancy and health impacts in the offspring. Pediatrics 133, e674–688.

Du, W. G., Zhao, B., Chen, Y. and Shine, R. (2011). Behavioral thermoregulation by turtle embryos. Proc Natl Acad Sci U S A 108, 9513–9515.

Duan, Y., Sun, Y. and Ambros, V. (2020). RNA-seq with RNase H-based ribosomal RNA depletion specifically designed for C. elegans. MicroPubl Biol 2020.

Elaswad, M. T., Watkins, B. M., Sharp, K. G., Munderloh, C. and Schisa, J. A. (2022). Large RNP granules in Caenorhabditis elegans oocytes have distinct phases of RNA-binding proteins. G3 (Bethesda) 12.

Ewels, P., Magnusson, M., Lundin, S. and Kaller, M. (2016). MultiQC: summarize analysis results for multiple tools and samples in a single report. Bioinformatics 32, 3047–3048.

Farley, B. M., Pagano, J. M. and Ryder, S. P. (2008). RNA target specificity of the embryonic cell fate determinant POS-1. RNA 14, 2685–2697.

Farley, B. M. and Ryder, S. P. (2008). Regulation of maternal mRNAs in early development. Crit Rev Biochem Mol Biol 43, 135–162.

Ghanta, K. S., Ishidate, T. and Mello, C. C. (2021). Microinjection for precision genome editing in Caenorhabditis elegans. STAR Protoc 2, 100748.

Ghanta, K. S. and Mello, C. C. (2020). Melting dsDNA Donor Molecules Greatly Improves Precision Genome Editing in Caenorhabditis elegans. Genetics 216, 643–650.

Hajdu-Cronin, Y. M., Chen, W. J. and Sternberg, P. W. (2004). The L-type cyclin CYL-1 and the heat-shock-factor HSF-1 are required for heat-shock-induced protein expression in Caenorhabditis elegans. Genetics 168, 1937–1949.

Hansen, P. J. (2009). Effects of heat stress on mammalian reproduction. Philosophical transactions of the Royal Society of London. Series B, Biological sciences 364, 3341–3350.

Holdorf, A. D., Higgins, D. P., Hart, A. C., Boag, P. R., Pazour, G. J., Walhout, A. J. M. and Walker, A. K. (2020). WormCat: An Online Tool for Annotation and Visualization of Caenorhabditis elegans Genome-Scale Data. Genetics 214, 279–294.

Huang, N. N., Mootz, D. E., Walhout, A. J., Vidal, M. and Hunter, C. P. (2002). MEX-3 interacting proteins link cell polarity to asymmetric gene expression in Caenorhabditis elegans. Development 129, 747–759.

Ilbay, O. and Ambros, V. (2019). Regulation of nuclear-cytoplasmic partitioning by the lin-28-lin-46 pathway reinforces microRNA repression of HBL-1 to confer robust cell-fate progression in C. elegans. Development 146.

Jones, A. R., Francis, R. and Schedl, T. (1996). GLD-1, a cytoplasmic protein essential for oocyte differentiation, shows stage- and sex-specific expression during Caenorhabditis elegans germline development. Developmental Biology 180, 165–183.

Jud, M. C., Czerwinski, M. J., Wood, M. P., Young, R. A., Gallo, C. M., Bickel, J. S., Petty, E. L., Mason, J. M., Little, B. A., Padilla, P. A., et al. (2008). Large P body-like RNPs form in C. elegans oocytes in response to arrested ovulation, heat shock, osmotic stress, and anoxia and are regulated by the major sperm protein pathway. Dev Biol 318, 38–51.

Jungkamp, A. C., Stoeckius, M., Mecenas, D., Grun, D., Mastrobuoni, G., Kempa, S. and Rajewsky, N. (2011). In vivo and transcriptome-wide identification of RNA binding protein target sites. Mol Cell 44, 828–840.

Kalchhauser, I., Farley, B. M., Pauli, S., Ryder, S. P. and Ciosk, R. (2011). FBF represses the Cip/Kip cell-cycle inhibitor CKI-2 to promote self-renewal of germline stem cells in C. elegans. EMBO J 30, 3823–3829.

Kawasaki, I., Shim, Y. H., Kirchner, J., Kaminker, J., Wood, W. B. and Strome, S. (1998). PGL-1, a predicted RNA-binding component of germ granules, is essential for fertility in C. elegans. Cell 94, 635–645.

Kaymak, E., Farley, B. M., Hay, S. A., Li, C., Ho, S., Hartman, D. J. and Ryder, S. P. (2016). Efficient generation of transgenic reporter strains and analysis of expression patterns in Caenorhabditis elegans using library MosSCI. Dev Dyn 245, 925–936.

Kurhanewicz, N. A., Dinwiddie, D., Bush, Z. D. and Libuda, D. E. (2020). Elevated Temperatures Cause Transposon-Associated DNA Damage in C. elegans Spermatocytes. Curr Biol 30, 5007–5017 e5004.

Li, R., Hu, K., Liu, H., Green, M. R. and Zhu, L. J. (2020). OneStopRNAseq: A Web Application for Comprehensive and Efficient Analyses of RNA-Seq Data. Genes (Basel) 11.

Love, M. I., Huber, W. and Anders, S. (2014). Moderated estimation of fold change and dispersion for RNA-seq data with DESeq2. Genome Biol 15, 550.

Lovegrove, B. G. (2014). Cool sperm: why some placental mammals have a scrotum. Journal of Evolutionary Biology 27, 801–814.

Merritt, C., Rasoloson, D., Ko, D. and Seydoux, G. (2008). 3’ UTRs are the primary regulators of gene expression in the C. elegans germline. Curr Biol 18, 1476–1482.

Miska, E. A., Alvarez-Saavedra, E., Abbott, A. L., Lau, N. C., Hellman, A. B., McGonagle, S. M., Bartel, D. P., Ambros, V. R. and Horvitz, H. (2007). Most Caenorhabditis elegans microRNAs are individually not essential for development or viability. PLoS Genet 3, e215.

Pagano, J. M., Farley, B. M., Essien, K. I. and Ryder, S. P. (2009). RNA recognition by the embryonic cell fate determinant and germline totipotency factor MEX-3. Proc Natl Acad Sci U S A 106, 20252–20257.

Riabinin, K., Kozhevnikov, M. and Ishay, J. S. (2004). Ventilating activity at the hornet nest entrance. Journal of Ethology 22, 49–53.

Rogers, A. K. and Phillips, C. M. (2020). RNAi pathways repress reprogramming of C. elegans germ cells during heat stress. Nucleic Acids Res 48, 4256–4273.

Ryder, S. P., Frater, L. A., Abramovitz, D. L., Goodwin, E. B. and Williamson, J. R. (2004). RNA target specificity of the STAR/GSG domain post-transcriptional regulatory protein GLD-1. Nat Struct Mol Biol 11, 20–28.

Sandberg, W. S., Schlunk, P. M., Zabin, H. B. and Terwilliger, T. C. (1995). Relationship between in vivo activity and in vitro measures of function and stability of a protein. Biochemistry 34, 11970–11978.

Schreiner, W. P., Pagliuso, D. C., Garrigues, J. M., Chen, J. S., Aalto, A. P. and Pasquinelli, A.E. (2019). Remodeling of the Caenorhabditis elegans non-coding RNA transcriptome by heat shock. Nucleic Acids Res 47, 9829–9841.

Spirin, A. S. (1966). “Masked” forms of mRNA. Curr Top Dev Biol 1, 1–38.

Stiernagle, T. (2006). Maintenance of C. elegans. WormBook : the online review of C. elegans biology, 1–11.

Surya, A., Bolton, B. M., Rothe, R., Mejia-Trujillo, R., Zhao, Q., Leonita, A., Liu, Y., Rangan, R., Gorusu, Y., Nguyen, P., et al. (2024). Cytosolic Ribosomal Protein Haploinsufficiency affects Mitochondrial Morphology and Respiration. bioRxiv.

Tabara, H., Hill, R. J., Mello, C. C., Priess, J. R. and Kohara, Y. (1999). pos-1 encodes a cytoplasmic zinc-finger protein essential for germline specification in C. elegans. Development 126, 1–11.

Tsukamoto, T., Gearhart, M. D., Spike, C. A., Huelgas-Morales, G., Mews, M., Boag, P. R., Beilharz, T. H. and Greenstein, D. (2017). LIN-41 and OMA Ribonucleoprotein Complexes Mediate a Translational Repression-to-Activation Switch Controlling Oocyte Meiotic Maturation and the Oocyte-to-Embryo Transition in Caenorhabditis elegans. Genetics 206, 2007–2039.

Walsh, B. S., Parratt, S. R., Hoffmann, A. A., Atkinson, D., Snook, R. R., Bretman, A. and Price, T. A. R. (2019). The Impact of Climate Change on Fertility. Trends Ecol Evol 34, 249–259.

Wang, J. T. and Seydoux, G. (2013). Germ cell specification. Adv Exp Med Biol 757, 17–39.

Wang, S., Ochoa, S. D., Khaliullin, R. N., Gerson-Gurwitz, A., Hendel, J. M., Zhao, Z., Biggs, R., Chisholm, A. D., Desai, A., Oegema, K., et al. (2019). A high-content imaging approach to profile C. elegans embryonic development. Development 146.

Wang, Y., Opperman, L., Wickens, M. and Hall, T. (2009). Structural basis for specific recognition of multiple mRNA targets by a PUF regulatory protein. Proc Natl Acad Sci U S A 106, 20186–20191.

Wright, J. E., Gaidatzis, D., Senften, M., Farley, B. M., Westhof, E., Ryder, S. P. and Ciosk, R. (2011). A quantitative RNA code for mRNA target selection by the germline fate determinant GLD-1. EMBO J 30, 533–545.

Zhang, B. L., Gallegos, M., Puoti, A., Durkin, E., Fields, S., Kimble, J. and Wickens, M. P. (1997). A conserved RNA-binding protein that regulates sexual fates in the C-elegans hermaphrodite germ line. Nature 390, 477-

