## Supplemental Table 1 for "The *mex*-3 3’ untranslated region is essential for reproduction during temperature stress"

Supplementary Table 2

| Primer Name | Purpose | Sequence |
| --- | --- | --- |
| mex-3_005F | PCR Confirmation of mex-3 3’ UTR genotypes | 5’ -CGATCATACTCTCGTGCCGA - 3’ |
| mex-3_007R | PCR Confirmation of mex-3 3’ UTR genotypes | 5’ -CTGAAACAATGGGACACCTCAA - 3’ |
| nos-2_H2B_01F | PCR Confirmtion of nos-2 marker | 5’ -TACACATGGCATGGATGAACT - 3’ |
| Nos-2_H2B_01R | PCR Confirmation of nos-2 marker | 5’ -AAGGCTATGAACGGGTAACTCA - 3’ |
| Linker1_pgl1_HA_F | Generation of pgl-1::mCherry reporter template | 5’ -TTAAATATTTATTTCAGTTTCATCCATTTCACATGTCCGGAGGGAGTGGA - 3’ |
| Linker2_pgl1_HA_R | Generation of pgl-1::mCherry reporter template | 5’ - CCACCGAAATCCACAATTTCTCGCTTGTTAGCCTCAGAACCTCCGCCACC - 3’ |
| CD.HC9.YJVG2684.AB | Guide RNA for the generation of spr20 | 5’ – GUUUCAUCCAUUUCACAUGG – 3’ |
| WRM81/82_Sequence_F1 | Sequencing and PCR confirmation of spr20 | 5’ – GAGTTTATGCGTTTCAAGGTG – 3’ |
| WRM81/82_Sequence_F2 | Sequencing and PCR confirmation of spr20 | 5’ – TCCACAGTTCATGTATGGAAG – 3’ |
| WRM81/82_Sequence_F3 | Sequencing and PCR confirmation of spr20 | 5’ – CTATGGGATGGGAAGCTTC – 3’ |
| WRM81/82_Sequence_R1 | Sequencing and PCR confirmation of spr20 | 5’ – GCCGGATGTTTAACATAAGC – 3’ |
| WRM81/82_Sequence_R2 | Sequencing and PCR confirmation of spr20 | 5’ – CCGTCTTCAGGGTACATTC – 3’ |
| WRM81/82_Sequence_R3 | Sequencing and PCR confirmation of spr20 | 5’ – CAATTCATCCATGCCACCT – 3’ |
