## Supplementary figures and images for "The *mex*-3 3’ untranslated region is essential for reproduction during temperature stress"

### Supplemental Figure 1

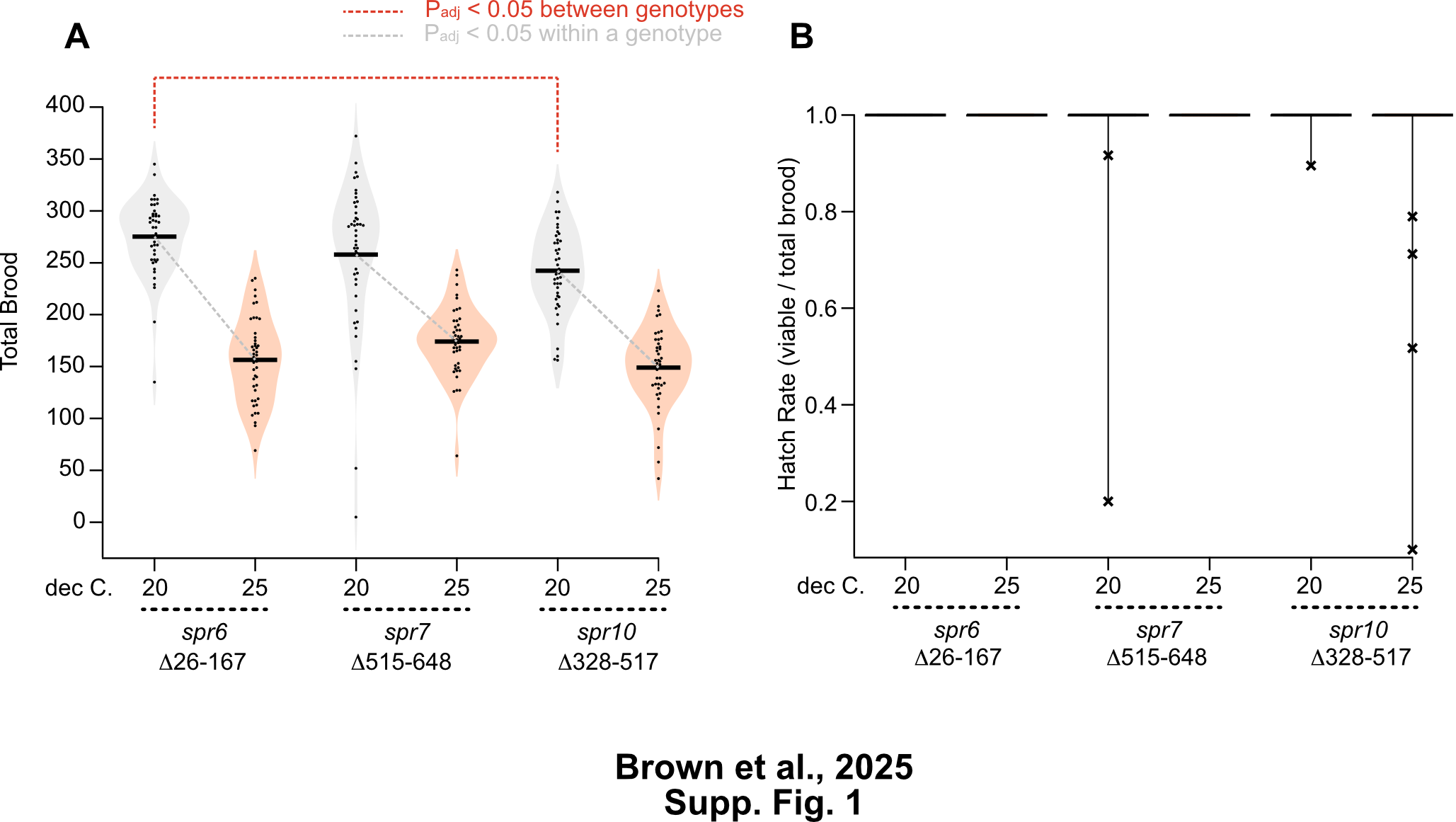

### Supplemental Figure 2

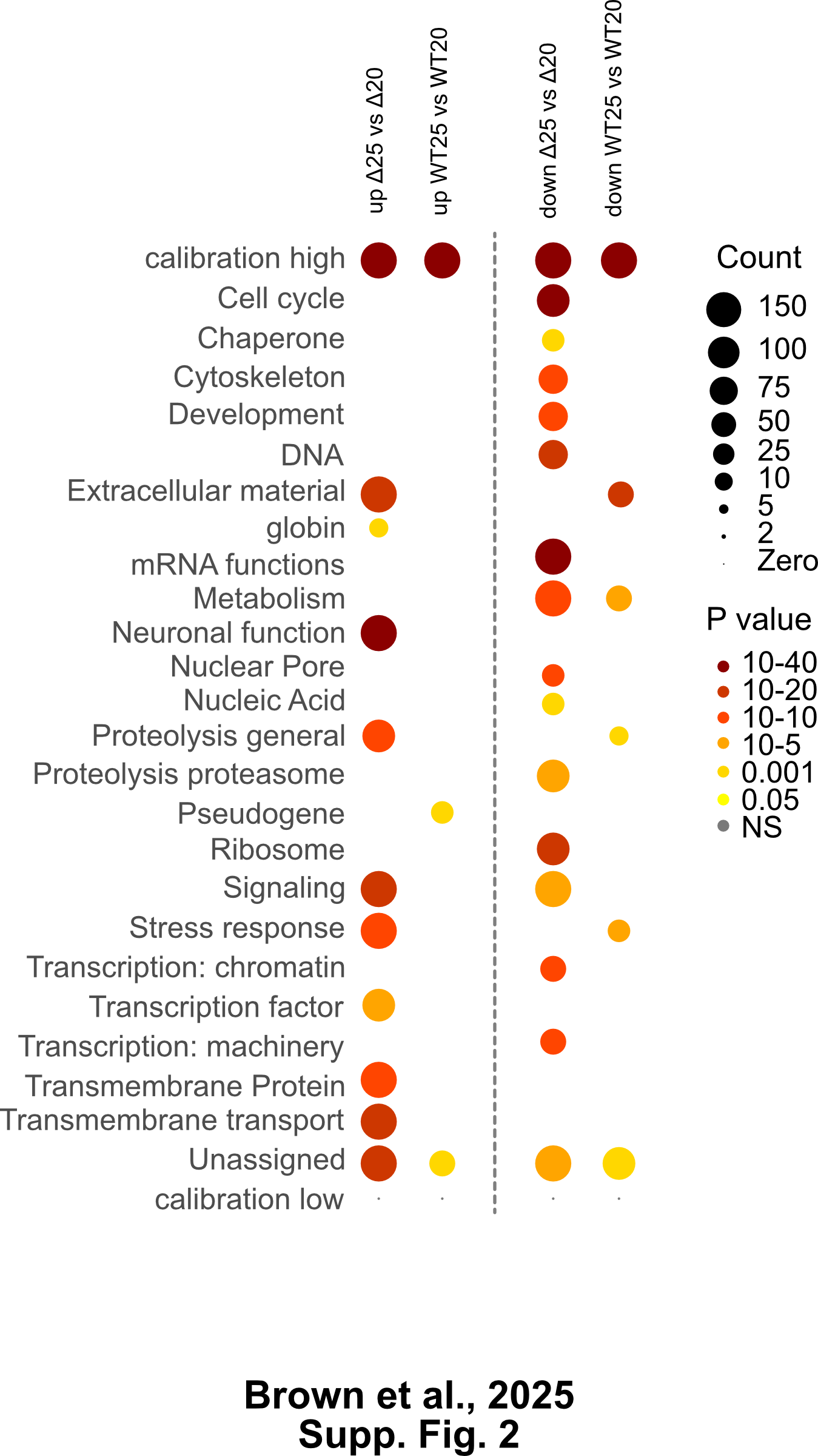
